# Coordinated cerebellar–midbrain activity for predictive signaling during fear learning in mice

**DOI:** 10.64898/2026.10.17.759883

**Authors:** Julie Danielle Urrutia Desmaison, Clément Léna, Daniela Popa

**Author notes:** These authors jointly supervised the work (Clément Léna, Daniela Popa). Corresponding authors: Clément Léna, & Daniela Popa.

## Abstract

Learning from experience to assess threat is crucial for survival and mental health. This function is known to involve several midbrain areas, notably the ventrolateral periaqueductal grey (vlPAG) and dorsal raphe nuclei (DRN). Recently it has been shown that fear learning involves projections from the cerebellar fastigial nucleus (FN) to the vlPAG. Here, we show using simultaneous single-unit recordings during fear conditioning that a subset of vlPAG/DRN and FN neurons acquire responses to conditioned cues following learning. Strikingly, these populations exhibit firing covariance and directional functional interactions consistent with their collaboration to the response to cues predicting threat. We also demonstrated the reciprocal monosynaptic connectivity between vlPAG/DRN and FN supporting the interactions between these structures. Together, our findings demonstrate shared dynamical activity within a network integrating midbrain and cerebellar computations during fear learning.

**IN BRIEF:** The projections of the cerebellar fastigial nucleus (FN) to the midbrain are involved in associative fear learning. Here, we show that during fear learning FN and periaqueductal grey/dorsal raphe (vlPAG/DRN) neurons acquire simultaneously responses to stimuli predicting aversive stimulus. Moreover, neurons encoding such predictive signals in the FN exhibit coordinated activity with the vlPAG/DRN, with directional functional interactions, suggesting that cerebellar – midbrain interactions contribute to the representation of aversive predictions during fear learning. We further characterize monosynaptic projections from the vlPAG/DRN to the contralateral FN, providing an anatomical substrate for reciprocal communications between the cerebellum and midbrain and extending previous descriptions of this pathway.

**HIGHLIGHTS:**

- Cerebellar fastigial (FN) neurons acquire stimulus specific predictive signals during fear learning.
- FN neurons encoding predictive signals exhibit coordinated activity and directional functional interactions with vlPAG/DRN neurons.
- vlPAG/DRN neurons projecting to the FN elicit short-latency responses in FN neurons.
- Reciprocal projections link the FN and vlPAG/DRN providing a potential substrate for cerebellar-midbrain interactions during fear learning.

## INTRODUCTION

Fear learning is a fundamental adaptive process that enables animals to recognize and respond to life-threatening situations, thereby ensuring survival^1,2^. Dysfunctions in the mechanisms underlying the acquisition or extinction of fear memories are central to several psychiatric disorders in humans, including post-traumatic stress disorder or generalized anxiety disorder, which severely affect quality of life. Understanding the neural mechanisms that support associative fear learning is therefore essential for advancing our knowledge of emotional processing and mental health. During associative fear learning, a neutral cue becomes predictive of an aversive event after the two have been repeatedly paired. This process, first described by Pavlov^3^, is driven by prediction error signals (the difference between expected and actual outcomes) which diminish as learning progresses^4,5^. Neural correlates of this associative process have been identified across multiple brain regions, with activity shifting from the responses to the aversive unconditioned stimulus (US) to the conditioned stimulus (CS) as learning occurs. This transition was initially observed in the lateral amygdala^6^ and subsequently in other regions such as the ventrolateral periaqueductal gray (vlPAG)^7,8^, establishing predictive neural signals bound to the CS as a hallmark of associative learning.

Fear learning arises from the coordinated activity of a distributed neuronal network that includes the amygdala, thalamus, hippocampus and prefrontal cortex^9,10^. The ventrolateral periaqueductal gray (vlPAG), initially thought as a region only mediating defensive behavior, has since been show to be part of the fear learning network, through its connections with forebrain regions as the amygdala and thalamus, as well as with the cerebellum^7,9,11^.

Neurons located in the border region of vlPAG and the neighboring dorsal raphe nuclei (DRN) acts as a multifunctional hub during fear learning. This vlPAG/DRN border region thus appears to serve as a multi-functional hub ^8,12,13^. It mediates behavioral and physiological responses such as freezing^10,14^ and pain modulation^15,16^. Neuronal activity in this area reflects threats^12,13,17–19^ and, during associative fear learning, exhibits a transfer of responses from aversive stimuli to cues predicting the occurrence of these aversive stimuli ^7,8^. Through connections with the amygdala and other forebrain regions^7,20,21^, the vlPAG/DRN therefore provides signals that shape the progression of fear learning.

Recent studies have revealed direct projections from the cerebellar fastigial nucleus (FN) to the vlPAG/DRN and have demonstrated their influence on aversive learning^22–28^. Indeed, emerging evidence implicates the cerebellum as a major contributor to emotional regulation and aversive learning^29–32^. Activity in the cerebellar hemispheres and vermis has been shown to reflect emotional valence and fear-related processes^31,33^, and cerebellar damage leads to affective instability and impaired behavioral modulation, core features of the cerebellar cognitive affective syndrome^34,35^. More specifically, inhibition or excitation of the monosynaptic projections from the FN to the vlPAG/DRN has been shown to modulate fear learning^23,25^. Moreover, stimulation of these projections at the time of prediction error signaling in the vlPAG/DRN was sufficient to alter fear learning, suggesting that these projections contribute to prediction-error signaling in vlPAG/DRN activity^7,8,36^. Interestingly, a previous report in cats suggested the existence of connections from the vlPAG/DRN back to the contralateral FN^37^, raising the possibility that reciprocal interactions between the cerebellum and vlPAG/DRN are involved in fear learning.

Reciprocal interactions between structures within the fear network are central to the adaptive encoding of threat. While FN and vlPAG/DRN are interconnected and have been recorded separately during fear learning^7,23,25^, the coordination of the activity between these regions has not been investigated. In this study, we aimed to characterize bidirectional interactions between the cerebellar fastigial nucleus and the midbrain vlPAG/DRN during fear learning to further understand the mechanisms underlying cerebellar modulation of fear learning. Specifically, we (i) identified neural populations in the FN and vlPAG/DRN that acquired predictive-signal responses to the conditioned stimulus during associative fear learning; (ii) examined the coupling between neural populations encoding predictive signal simultaneously in both structures to study their coordinated contributions to fear learning and (iii) identified and mapped anatomical projections from the vlPAG/DRN to the FN suggesting reciprocal communication between these regions.

## RESULTS

### FN and vlPAG/DRN neural population responses are altered by fear conditioning

To characterize neural dynamics encoding fear-stimulus associations in the FN and the vlPAG/DRN border region, we monitored neural activity through simultaneous extracellular recordings from permanently implanted electrodes in both structures in freely moving mice during a CS+/CS- Pavlovian fear conditioning paradigm (Fig.1 a, e). During the fear conditioning paradigm, animals were presented with an equal number of acoustic CS+ and CS- stimuli, consisting of either a 30-second pure tone at 7000Hz or white noise, counterbalanced across animals. The CS+ always co-terminated in a mild aversive foot shock (US), while the CS- was never paired with the US (Fig.1 b). Fear learning success was monitored by measuring the animal’s immobility levels during the CS presentation. Immobility increased from comparable levels during CS+ and CS- presentation during habituation to significantly higher levels during CS+ presentation during recall on the following day (Fig.1 f). However, the relatively high immobility observed in response to CS- suggested the presence of a mild fear generalization. All animals (n=12) were habituated to the acoustic CS+ and CS- stimuli prior to fear conditioning (Fig.1 f).

**Figure 1.**
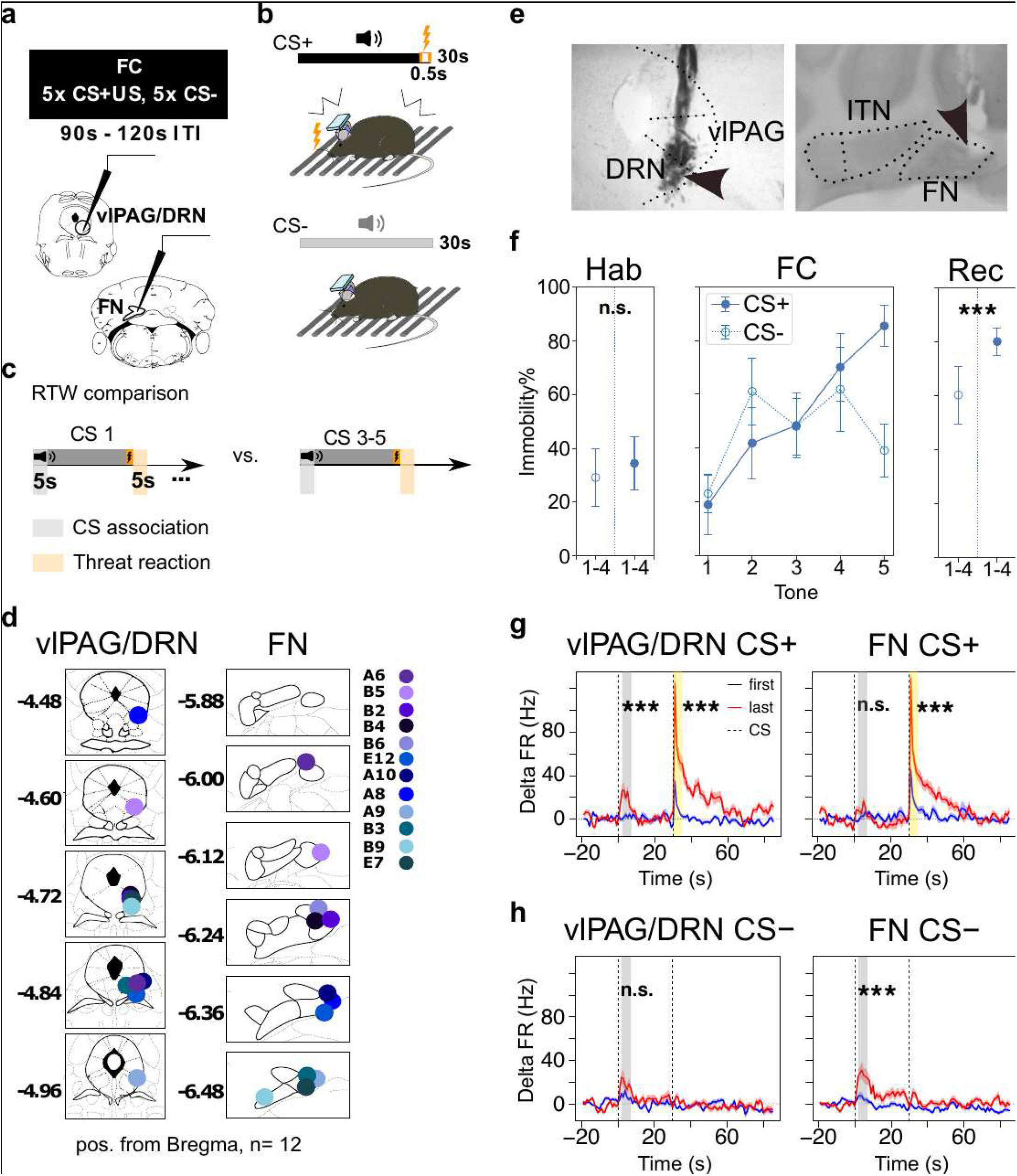
FN and vlPAG/DRN neural population response is altered by fear conditioning. (a) Fear conditioning paradigm with bundle recordings from vlPAG/DRN and FN. (b) CS+ coterminating with a footshock, CS- without footshock. (c) Definition of reporter time windows (RTW) (d) Electrode insertion sites in FN and vlPAG/DRN, color legend identifies animal. (e) Electrode insertion sites. (f) Immobility levels during habituation and fear conditioning. (g) Mean firing rate (FR) of all recorded units during CS+ and US, comparing RTW of first to last 3 presentations. (h) Mean firing rate (FR) of all recorded units during CS-, comparing RTW of first to last 3 presentations.

To characterize the simultaneous representation of fear-stimulus associations in the neural dynamics of the vlPAG/DRN and FN, we isolated units from our electrophysiological recordings and compared neural responses following CS onsets and US delivery. To this aim, we defined 5-second reporter time windows (RTWs) at the beginning of the CS and following the US, and compared responses to the first stimulus presentation, before any association has occurred, with the average response across last three presentations, when the association becomes established (Fig.1 c).

We isolated units from the right caudal vlPAG/DRN border region (n=88) and the contralateral medial FN (n=85) (Fig.1 d) using electrode bundles emerging from guide cannulas implanted just above the targeted structures (Fig.1 f).

At a population level, neurons in vlPAG/DRN acquired a significant response to the CS+ onset (RTW CS+ first vs. last 3 vlPAG/DRN: p-value 6.99e-7) whereas no significant modulation was observed at CS- onset (RTW CS- first vs. last 3 vlPAG/DRN: p-value 0.057) following fear conditioning. This suggests that the association between CS+ and US resulted in the acquisition of a response to CS+ broadly in the vlPAG/DRN (Fig.1 g).

In contrast, no significant response at CS+ onset was observed at the population level in FN (RTW CS+ first vs. last 3 FN: p-value 0.181), whereas a significant response to CS- onset was observed (RTW CS- first vs. last 3 FN: p-value 0.006, Fig.1 h). These findings suggest that both FN and vlPAG/DRN adapt their neural activity patterns during fear conditioning, with increased responses to the aversive US and distinct patterns of adaptation to the CS+.

### CS+ specific predictive signals in FN and vlPAG/DRN subpopulations

The analysis above only considered the general behavior of the whole population; however, since the FN and vlPAG/DRN regions may be heterogeneous, we then examined whether subsets of neurons in each region exhibited more specific responses to CS+ onset following fear conditioning. Since the response to CS+ may reflect a change of threat expectation, i.e. increased prediction of an aversive event, we refer to these responses to the CS+ as ‘predictive signals’ (PSs). To identify acquired responses to CS+ onset, we calculated the change in firing rate between the first (naïve) CS+ presentation and the last 3 (trained) CS+ presentations for all units in both FN and vlPAG/DRN (Fig. 2a, c left panel). We then performed k-means clustering on the profile of firing rate changes within the CS+ RTW, and identified groups of units that either exhibited increased responses at CS+ onset (‘PS units’) or did not exhibit such changes (‘non-PS units’) in both the vlPAG/DRN and FN (Fig. 2a, c right panel. PS units, which showed a significant increase in firing rate at CS+ onset relative to their baseline state, were identified in both FN and vlPAG/DRN. Therefore, while FN did not show responses to CS+ at the population level, some units nonetheless exhibited such responses. The increase in firing rate at CS+ onset was significantly larger in PS units than in non-PS units in both the FN and vlPAG/DRN, confirming the relevance of the clustering approach for identifying acquired responses to CS+ in both structures (Fig. 2f, h).

**Figure 2.**
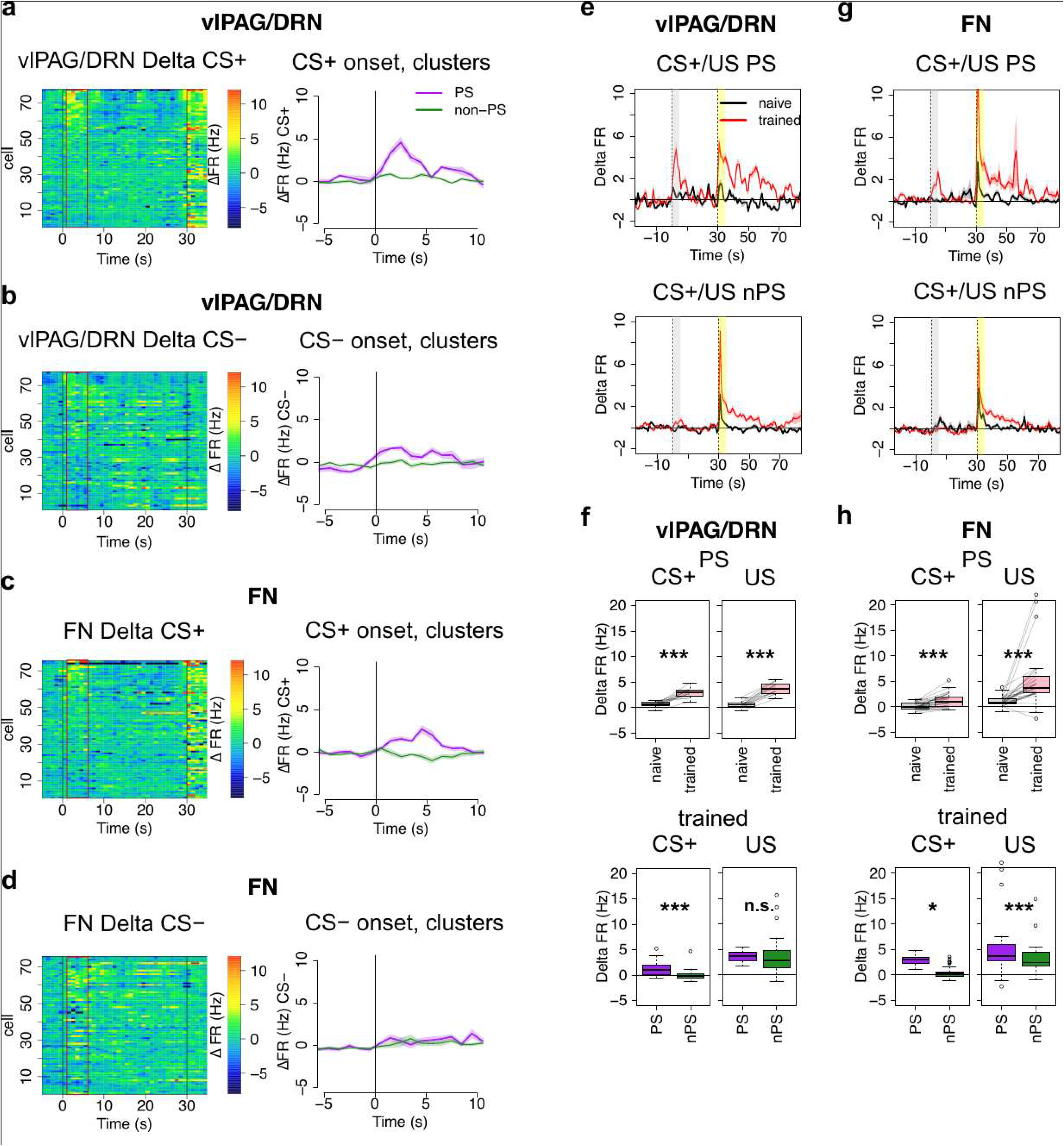
CS+ specific predictive signal in FN and vlPAG/DRN subpopulations. (a) Heatmap of delta FR CS + in vlPAG/DRN single neurons, CS+ onset at 0 (average last 3 - first), RTW in red rectangle. Mean +-SEM of delta FR in PS and non-PS cluster units in vlPAG/DRN. (b) Heatmap of delta FR CS - in vlPAG/DRN single neurons, CS- onset at 0 (average last 3 - first), RTW in red rectangle. Mean +-SEM of delta FR in PS and non-PS cluster units in vlPAG/DRN. (c) Heatmap of delta FR CS + in FN single neurons, CS+ onset at 0 (average last 3 - first), RTW in red rectangle. Mean +-SEM of delta FR in PS and non-PS cluster units in FN. (d) Heatmap of delta FR CS + in FN single neurons, CS+ onset at 0 (average last 3 - first), RTW in red rectangle. Mean +-SEM of delta FR in PS and non-PS cluster units in FN. (e) Mean +-SEM of FR during CS+ and US in PS and non-PS units in vlPAG/DRN, comparing naïve (first) to trained (average last 3) neural responses. (f) Quantification of RTW neural responses CS+ and US in PS and non-PS units in vlPAG/DRN, comparing naïve (first) to trained (average last 3) and trained PS to trained non-PS. (g) Mean +-SEM of FR during CS+ and US in PS and non-PS units in FN, comparing naïve (first) to trained (average last 3) neural responses. (h) Quantification of RTW neural responses CS+ and US in PS and non-PS units in FN, comparing naïve (first) to trained (average last 3) and trained PS to trained non-PS.

**Figure 2 Supplementary Fig. 1:**
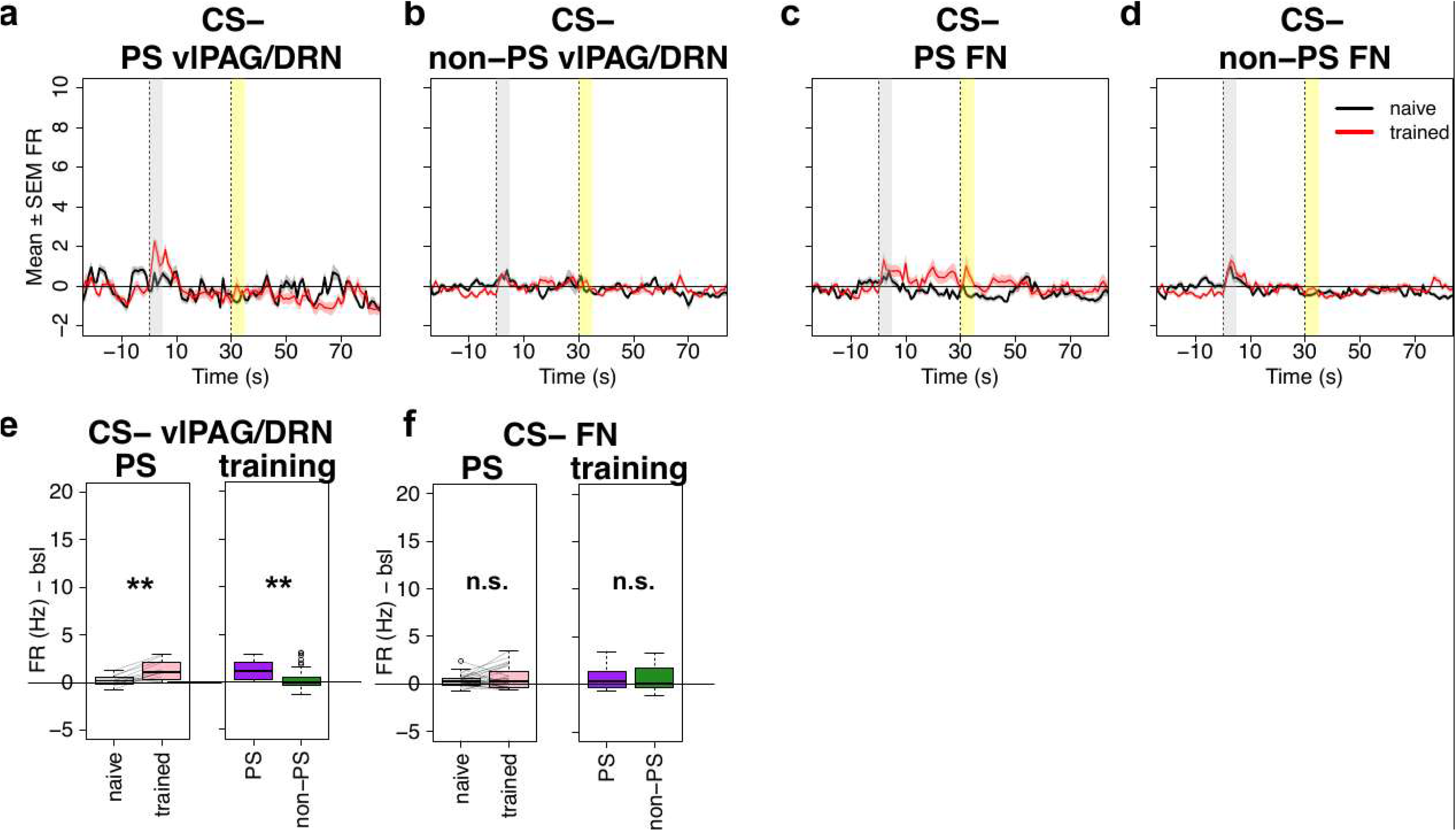
Population neural responses to CS- in vlPAG/DRN in PS and non-PS cells. (a) PS neurons: Mean FR in Hz comparing trained (red) to untrained neural response (black) during CS- in vlPAG/DRN (b) non-PS neurons: Mean FR in Hz comparing trained (red) to untrained neural response (black) during CS- in vlPAG/DRN (c) PS neurons: Mean FR in Hz comparing trained (red) to untrained neural response (black) during CS- in FN (d) non-PS neurons: Mean FR in Hz comparing trained (red) to untrained neural response (black) during CS- in FN (e) Quantification Mean FR in Hz during reporter time windows comparing trained (red) to untrained neural response (black) during CS- and trained PS to trained non-PS in vlPAG/DRN (f) Quantification Mean FR in Hz during reporter time windows comparing trained (red) to untrained neural response (black) during CS- and trained PS to trained non-PS in FN

To confirm that PS and non-PS clusters represented specific acquisition of CS+ onset responses, we calculated changes in firing rate for all units between naïve and trained CS- presentations (Fig. 2c, d left panel, Fig 2S1). We did not observe a general acquired response to CS-onset in either group, confirming that the PS cluster represents units that specifically acquired a response to CS+ onset in both FN and vlPAG/DRN (Fig. 2c, d right panel). Interestingly, PS units in both the FN and vlPAG/DRN also exhibited a pronounced increase in their response to the US over the course of learning (Fig. 2e, g, f, h). Thus, subgroups of neurons in both FN and vlPAG/DRN acquired a stimulus specific predictive response accompanied by an increased response to the US following fear conditioning.

The PS neurons identified in the vlPAG/DRN are unlikely to mediate a purely aversive stimulus response, as they did not exhibit a strong response to the US prior to fear conditioning (Fig. 2 e). In contrast, non-PS neurons exhibited a clear response to the US prior to fear conditioning, suggesting that this population may include neurons involved in encoding aversive behavior and responses mediated by the vlPAG/DRN.

### Neural discharge of PS units in the FN and vlPAG/DRN is correlated and suggests bidirectional interactions

To examine whether FN and vlPAG/DRN populations exhibit tight temporal functional coordination, we examined the relation between their firing activity before and during fear learning.

We computed cross-correlograms between the spike trains of PS units recorded simultaneously in FN and vlPAG/DRN during the minutes preceding the fear conditioning paradigm and during fear conditioning (Fig.3 a, Fig. 3S1 a, b). Cross-correlograms were always computed with the FN unit as a reference (Fig.3 a). We used the shape of the cross-correlogram to determine the strength and temporal structure of correlations between unit pairs. Many unit pairs exhibited a central peak of a few milliseconds wide, indicating short time-scale coordination of discharge between FN and vlPAG/DRN PS neurons pairs as well as between non-PS neurons pairs already in baseline conditions. Some FN PS neurons also exhibited correlations with vlPAG/DRN non-PS neurons (Fig. 3S1 a). These data indicate that PS-FN neurons are more related functionally to vlPAG/DRN neurons than other FN neurons. Moreover, pairs of PS neurons appeared more likely to exhibit significant coupling than non-PS pairs (Fig. 3S1) suggesting that these neurons may belong to functionally related subnetworks.

**Figure 3.**
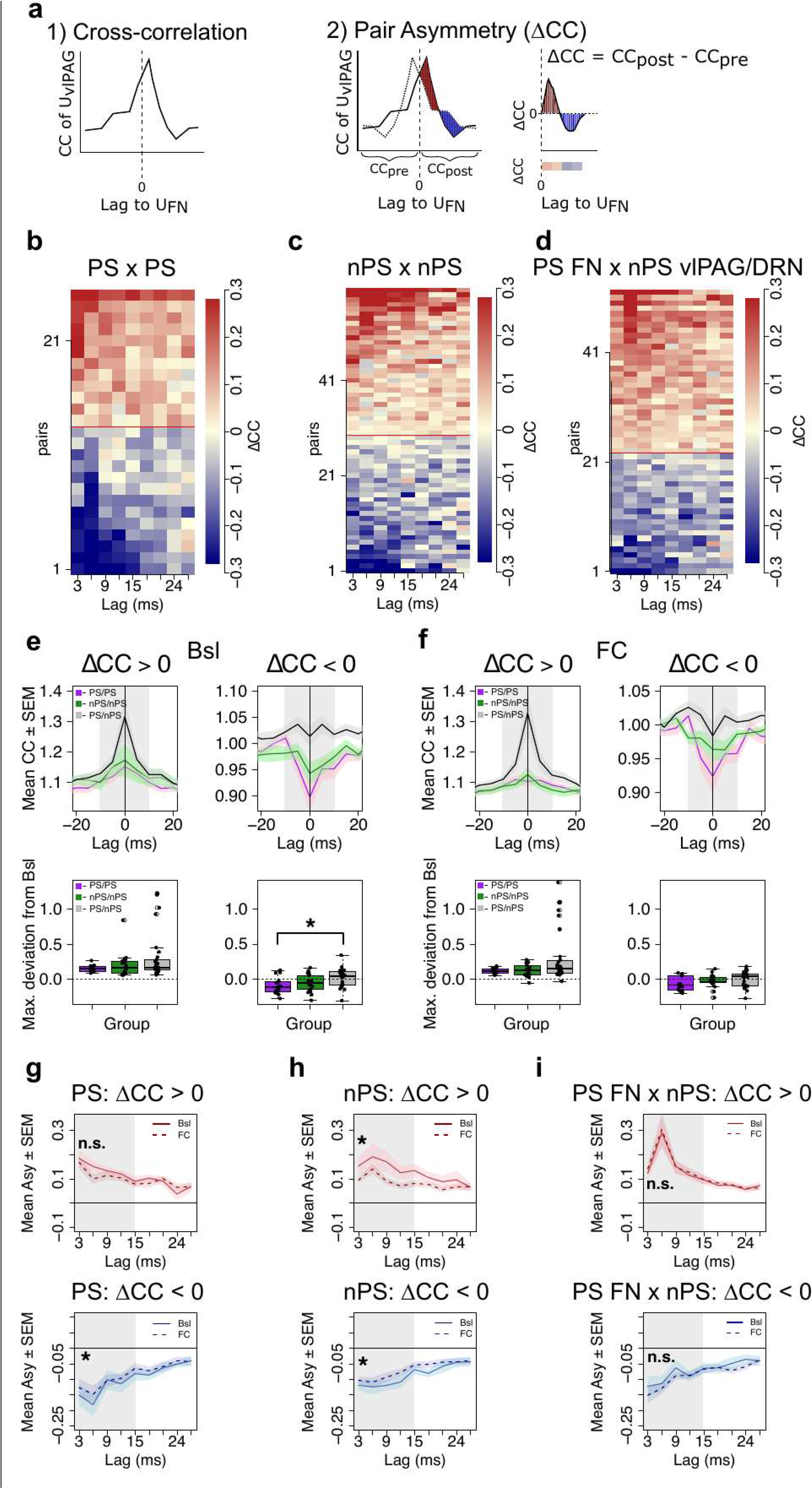
Evolution of FN and vlPAG/DRN bidirectional neural coupling over baseline behavior and fear conditioning. (a) Schematic representation of cross-correlation and pair asymmetry calculation. FN units are always treated as reference units. The red shaded area represents positive asymmetry, the blue shaded area represents negative asymmetry. (b) Asymmetry values obtained for correlated PS neuron pairs between vlPAG/DRN and FN during baseline conditions. (c) Asymmetry values obtained for correlated non-PS neuron pairs between vlPAG/DRN and FN during baseline conditions. (d) Asymmetry values obtained for correlated FN-PS and vlPAG/DRN non-PS neuron pairs between vlPAG/DRN and FN during baseline conditions. (e) Upper panel: Group mean cross-correlogram shapes for PSxPS, non-PS x non-PS and FN PS x vlPAG/DRN non-PS pairs during baseline conditions, grouped by positive and negative asymmetry. Lower panel: cross-correlogram shape extrema over 15 ms around center peak (grey area). (f) Upper panel: Group mean cross-correlogram shapes for PSxPS, non-PS x non-PS and FN PS x vlPAG/DRN non-PS pairs during fear conditioning, grouped by positive and negative asymmetry. Lower panel: cross-correlogram shape extrema over 15 ms around center peak (grey area). (g) PS pairs grouped by positive and negative Asymmetry and their coupling evolution over fear conditioning. (h) non-PS pairs grouped by positive and negative Asymmetry and their coupling evolution over fear conditioning. (i) FN PS x vlPAG/DRN non-PS pairs grouped by positive and negative Asymmetry and their coupling evolution over fear conditioning.

**Figure 3 Supplementary Fig. 1:**
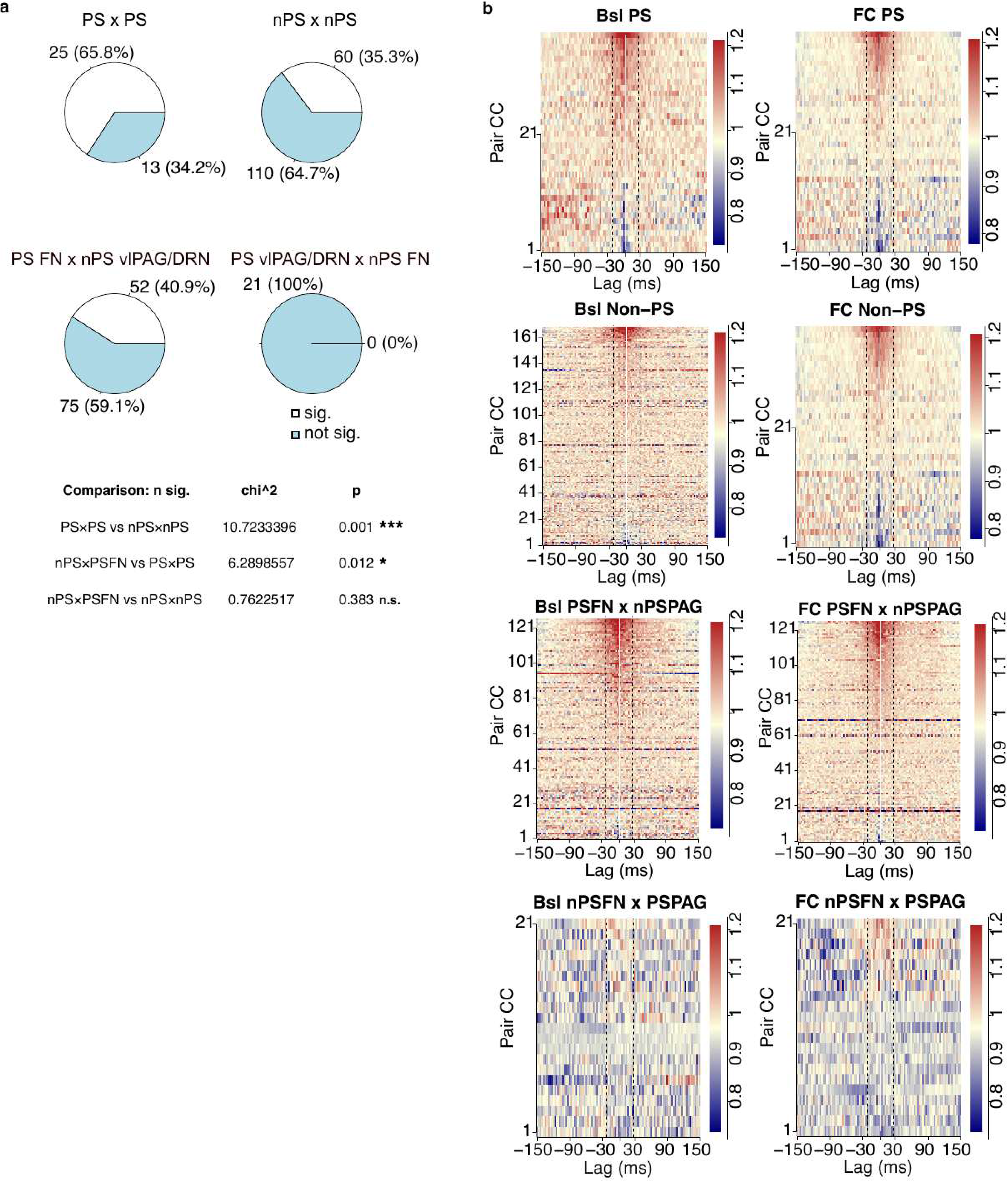
heatmaps of cross-correlograms. (a) Number of pairs detected between PS and non-PS subgroups of vlPAG/DRN and FN. Table below: chi^2 test comparing the proportion of significant neurons between subgroups (b) Baseline behavior PS and non-PS heatmaps of all pairings cross-correlations during baseline and fear conditioning

Moreover, we observed that a number of crosscorrelograms exhibited some degree of asymmetry, with an excess of vlPAG/DRN spikes either following or preceding FN discharge (‘positive’ vs ‘negative’ asymmetry, respectively). To quantify crosscorrelogram asymmetry, we subtracted each crosscorrelogram from its mirror image along the vertical axis. This difference in delayed coincidence between negative and positive time lags was denoted as **Δ**CC (Fig. 3 a). Pairs with an excess of vlPAG/DRN spikes following FN unit firing (positive **Δ**CC values for delays in the [3,30) ms interval) were considered as ‘positive asymmetry’ pairs, whereas pairs with an excess of vlPAG/DRN firing preceding FN firing (negative **Δ**CC values for delays in the [3,30) interval) were considered ‘negative asymmetry’ pairs. Significantly asymmetric pairs were identified both before and during fear conditioning among PS x PS pairs, non-PS x non-PS pairs and PS x non-PS pairs (Fig.3 b-d).

Among positive asymmetry pairs, the cross-correlogram dominantly exhibited central positive peaks, in line with shared inputs and/or excitatory interactions (Fig. 3e). This pattern remained stable under both baseline conditions and fear conditioning (Fig. 3 e,f). In contrast, negative asymmetry pairs rather exhibited cross-correlogram shapes with a central negative peak, more consistent with inhibitory interactions (Fig. 3e). Negative asymmetry pairs in both PS x PS pairs and non-PS x non-PS pairs consistently exhibited negative peaks, whereas FN PS x vlPAG/DRN non-PS pairs showed a mixed pattern (Fig. 3e). Interestingly, this distribution shifted toward negative peaks during fear conditioning (Fig. 3f). These findings suggest that inhibitory coordination patterns may contribute to the adaptation of neural responses to aversive conditions.

Positive asymmetry among PS-PS pairs was preserved between baseline and fear conditioning (Fig. 3g), while both positive and negative asymmetry decreased during fear conditioning in non-PS – non-PS pairs compared with baseline (Fig. 3 h). Interestingly, among FN PS neurons paired with vlPAG/DRN non-PS neurons, both negative and positive asymmetry remained stable from baseline to fear conditioning, while positive asymmetry remained prominent. This suggests that, in PS-PS pairs, the temporal relationship between FN and vlPAG/DRN discharge was maintained, while functional coordination among non-PS pairs decreased during fear conditioning. The preferred temporal order of firing was maintained between FN and vlPAG/DRN PS units, with FN discharge preceding vlPAG/DRN discharge, as well as in pairs in which FN PS units interacted with vlPAG/DRN non-PS neurons. Together, these findings suggest that FN PS neuron discharge are more strongly tuned on vlPAG/DRN activity during fear conditioning and in baseline conditions. They also raise the possibility that vlPAG/DRN non-PS neurons comprise functional subpopulations that become increasingly relevant for learning during fear conditioning, even though they do not initially exhibit the characteristic predictive signal response.

These findings indicate that vlPAG/DRN and FN units participate in shared functional networks exhibiting tight coordination of discharge patterns both before and during fear learning. Moreover, fear learning affects this coordination, notably by weakening the sequential activations of FN and vlPAG/DRN neurons within non-PS populations.

### vlPAG/DRN units with broader spikes preferentially respond to CS+ after fear conditioning

We then aimed to determine whether different neuronal subtypes could be distinguished in vlPAG/DRN based on their waveform characteristics. For each unit, we extracted recovery time, peak to trough width and amplitude ratio (trough-peak/(trough+peak)). We then performed k-means clustering after determining the optimal number of clusters using the silhouette criterion to identify different waveform types (Fig. 3S2 a).

We identified two clusters in vlPAG/DRN, C0 (recovery time 1.705 ± 0.041 ms SEM, spike half width 0.431 ± 0.013 ms SEM, amplitude ratio 0.386 ± 0.016 SEM) and C1 (recovery time 1.535 ± 0.036 ms SEM, spike half width 0.267 ± 0.009 ms SEM, amplitude ratio 0.512 ± 0.0137 SEM) (Fig. 3S2 b). The smaller amplitude ratio and large half width of vlPAG/DRN C0 neurons are consistent with waveform characteristics previously described for putative dopaminergic neurons in the VTA ^38^. The C0 waveform cluster contained a greater population of PS units in the vlPAG/DRN (n=9/13, Fig. 3S2 c). This distinction was also reflected in the response dynamics of the two populations of neurons, where C0 units developed a strong response to CS+- onset, but not to CS- onset over the course of fear conditioning, while C1 units developed only a weak response to both types of CS (Fig. 3S2 d). Thus, PS and non-PS neurons in the vlPAG/DRN may comprise distinct neuronal populations, with PS neurons being more likely to exhibit broader waveforms in association with the acquisition of a CS+ specific response.

**Figure 3 Supplementary Fig. 2:**
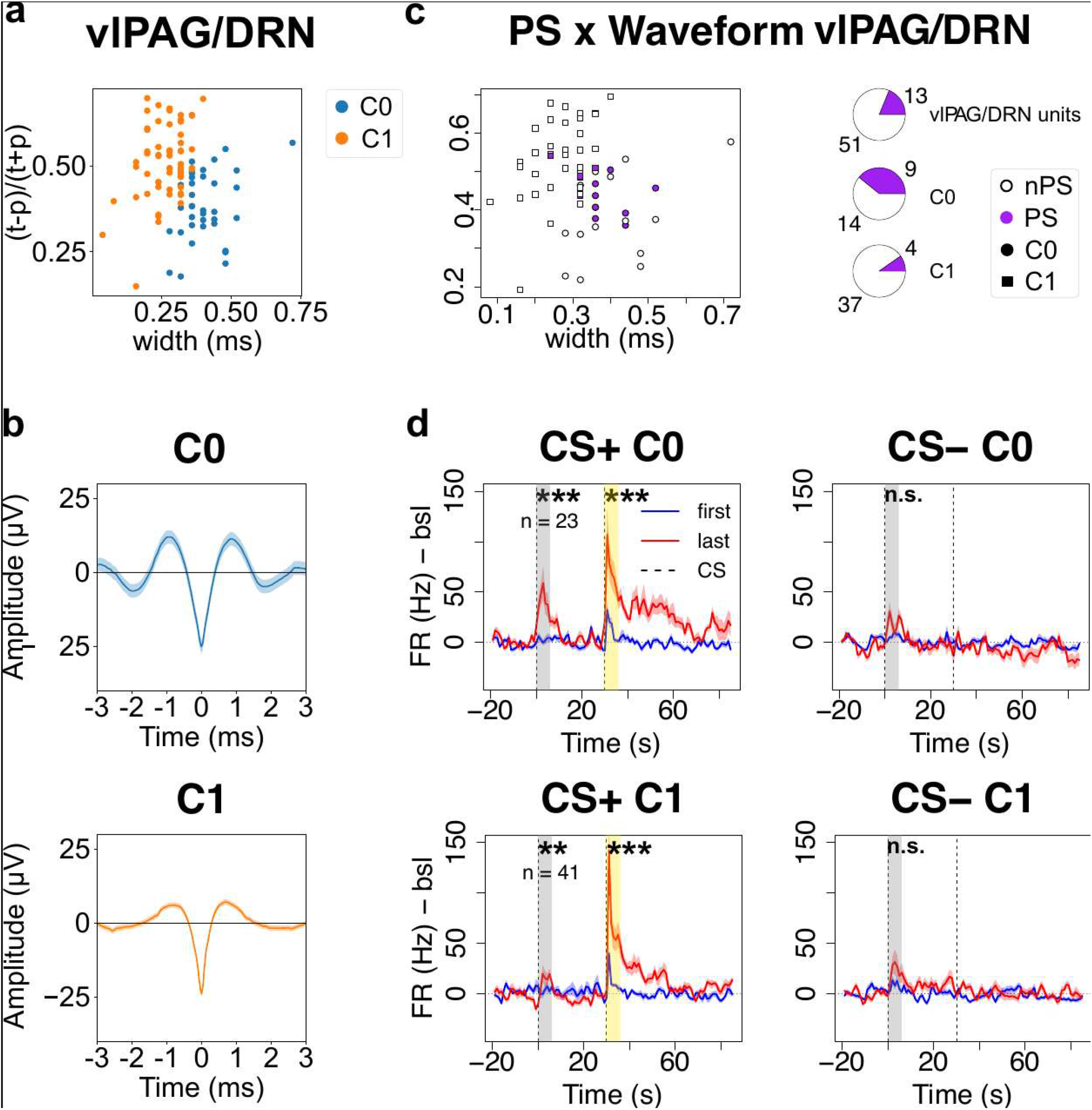
Waveform based clustering of vlPAG/DRN units. (a) Amplitude ratio against halfwidth of recorded waveforms according to cluster in vlPAG/DRN (b) Mean + SEM of cluster waveforms in vlPAG/DRN (c) Overlay of waveform clusters and PS or non-PS criterion in vlPAG/DRN units on the left, quantification of PS units per cluster and in all DRN units on the right. (d) Mean FR of naïve and trained waveform clusters and quantification of FR differences in reporter time windows: PAG C0 units: response to US, p= 4.8e-8, to CS+ p=1.3e-5, to CS- p=0.32. PAG C1 units response to US: p=9.7e-10, to CS+: p=5.9e-3, to CS-: p= 0.12.

### Monosynaptic projections from vlPAG/DRN to FN

The short-term sequential activations from FN and vlPAG/DRN neurons, evidenced above by ‘positive asymmetry’ in the crosscorrelograms, could involve the excitatory projections from the FN to the vlPAG/DRN area^23^. To further investigate the anatomical substrates that could underlie the ‘negative asymmetry’ observed in some pairs (i.e. FN neurons preferentially discharging after vlPAG/DRN), we examined whether the vlPAG/DRN reciprocally projects to the FN. To date, such monosynaptic projections from the vlPAG/DRN to the contralateral FN nucleus have only been described in cats^37^, and we therefore aimed to determine whether they are also present in mice.

To demonstrate the existence of vlPAG/DRN neurons projecting to the FN, we performed retrograde labelling from the FN by combining unilateral injections of a retrograde AAVretro-mCherry-cre virus in the FN with an injection of a Cre-dependent AAV-pCAG-FLEX-eGFP virus in the contralateral vlPAG/DRN of 5 male C57Bl6N mice (Fig.4 a). This approach revealed distinct labeled somata in the posterior ventrolateral PAG and adjacent lateral DRN with fewer labeled neurons in the laterodorsal tegmental nucleus (LDT), confirming the existence of direct projections from the vlPAG/DRN to the FN (Fig.4 b, d).

**Figure 4.**
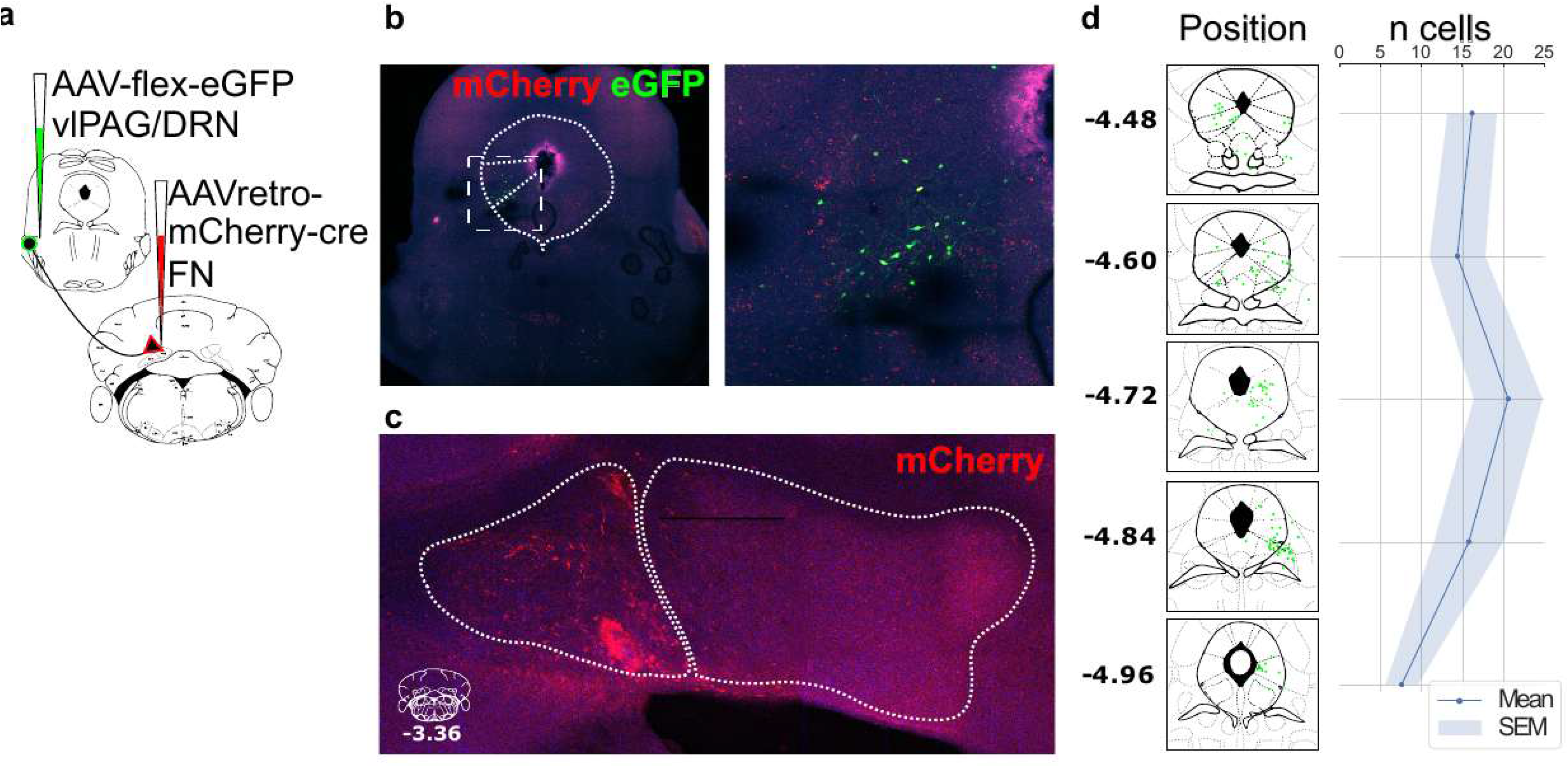
monosynaptic projections from the vlPAG/DRN to FN. (a) Schematic representation of the viral tracing strategy: injection of AAV-flex-eGFP in vlPAG/DRN and retrograde AAV-mCherry-cre in FN. (b) Expression of eGFP and mCherry in the midbrain and zoom-in on the vlPAG/DRN, (c) Expression of mCherry in the cerebellar nuclei and zoom-in on the injection site in the FN. (d) Example of distribution from one animal (left) and quantification for all animals (right, n=5) of eGFP-expressing cell bodies in coronal sections of vlPAG/DRN.

### vlPAG/DRN optogenetic stimulation induces short-latency responses in the FN

To assess the effect of direct vlPAG/DRN inputs on FN neuronal discharge, we performed extracellular electrophysiological recordings in the FN in mice expressing channel-rhodopsin-2 (ChR2) specifically in neurons projecting to the FN (Fig. 5S1 a, c) following retrograde rAAV-Chr2-eYFP injections in the FN (Fig. 5 a, 5S1 a). Optogenetic stimulation of these neurons in the vlPAG/DRN via an implanted optic fiber (Fig. 5S1 b) in awake, freely moving mice elicited robust and rapid (<10ms) negative LFP deflections in the FN, with response amplitudes scaling with light intensity (Fig.5 b). Increased firing of FN neurons in response to vlPAG/DRN stimulation (Fig. 5 c, e) was observed in 13 neurons from 3 mice. Response latencies ranged from 15 to 60 ms, consistent with both direct and polysynaptic activation pathways (Fig. 5 b, c). The average modulation of firing rate in responsive units during stimulation also increased with light intensity (Fig. 5 c). These findings indicate that vlPAG/DRN neurons projecting to the FN can exert a strong and rapid excitatory influence on FN neurons. Together with the anatomical evidence for monosynaptic vlPAG/DRN to FN projections, these results support the existence of a functional circuit linking between vlPAG/DRN and FN neuronal populations.

**Figure 5.**
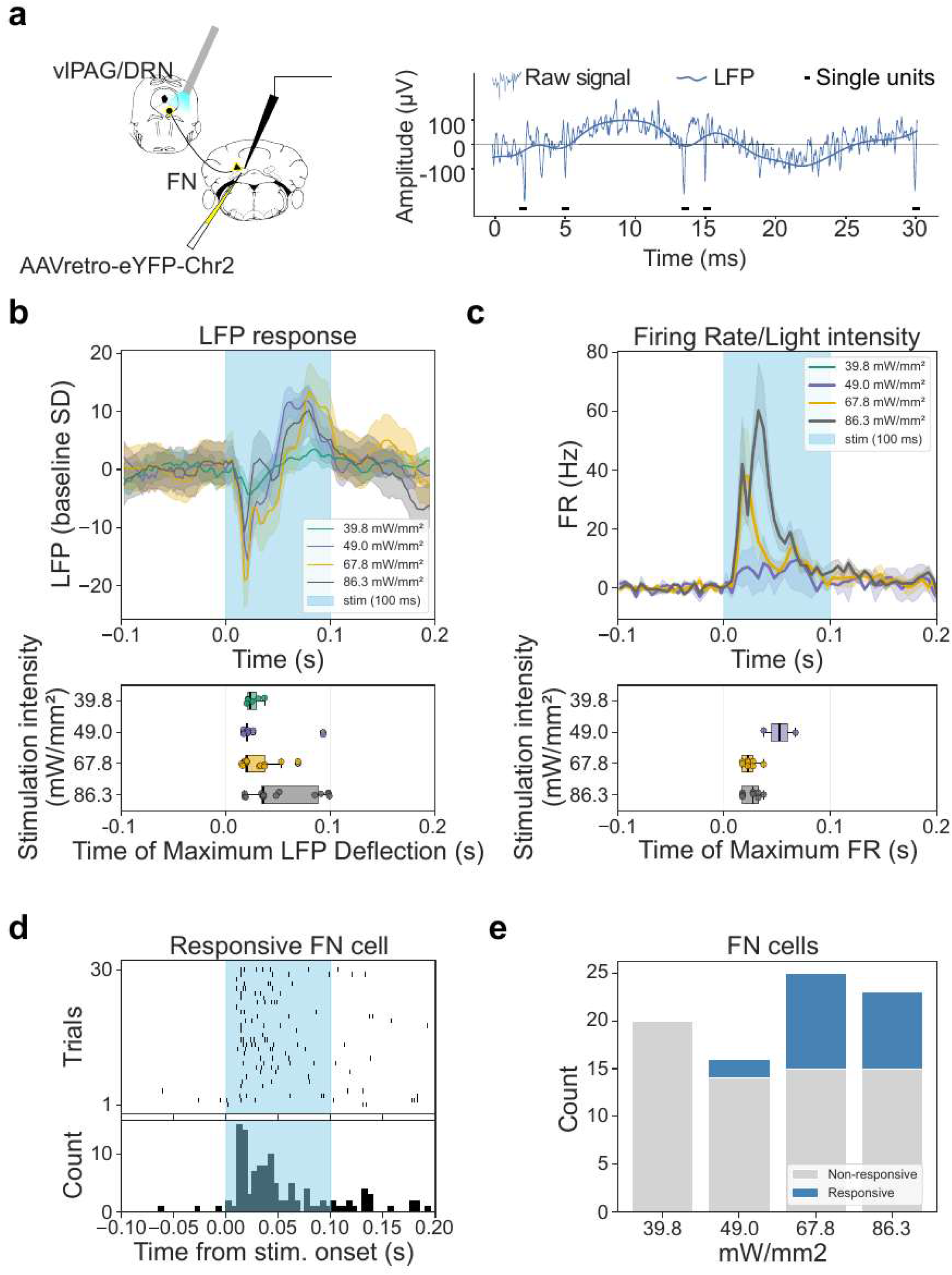
Laser stimulation of Chr2-expressing neurons in vlPAG/DRN and extracellular recordings in FN. (a) Experimental setup and processing of extracellular recording signal, spike and LFP extraction. (b) Upper panel: LFP response in FN to laser stimulation in vlPAG/DRN, SEM in shaded area. Lower panel: time of maximum deflection in single channels. (c) Upper panel: Average FR during laser stimulation in responsive cells in FN, SEM in shaded area. Lower panel: Time of maximum FR in single units. (d) Example of FN unit responding to vlPAG/DRN optogenetic stimulation. (e) Number of responsive and non-responsive units in FN, in 3 animals.

**Figure 5 Supplementary Fig. 1:**
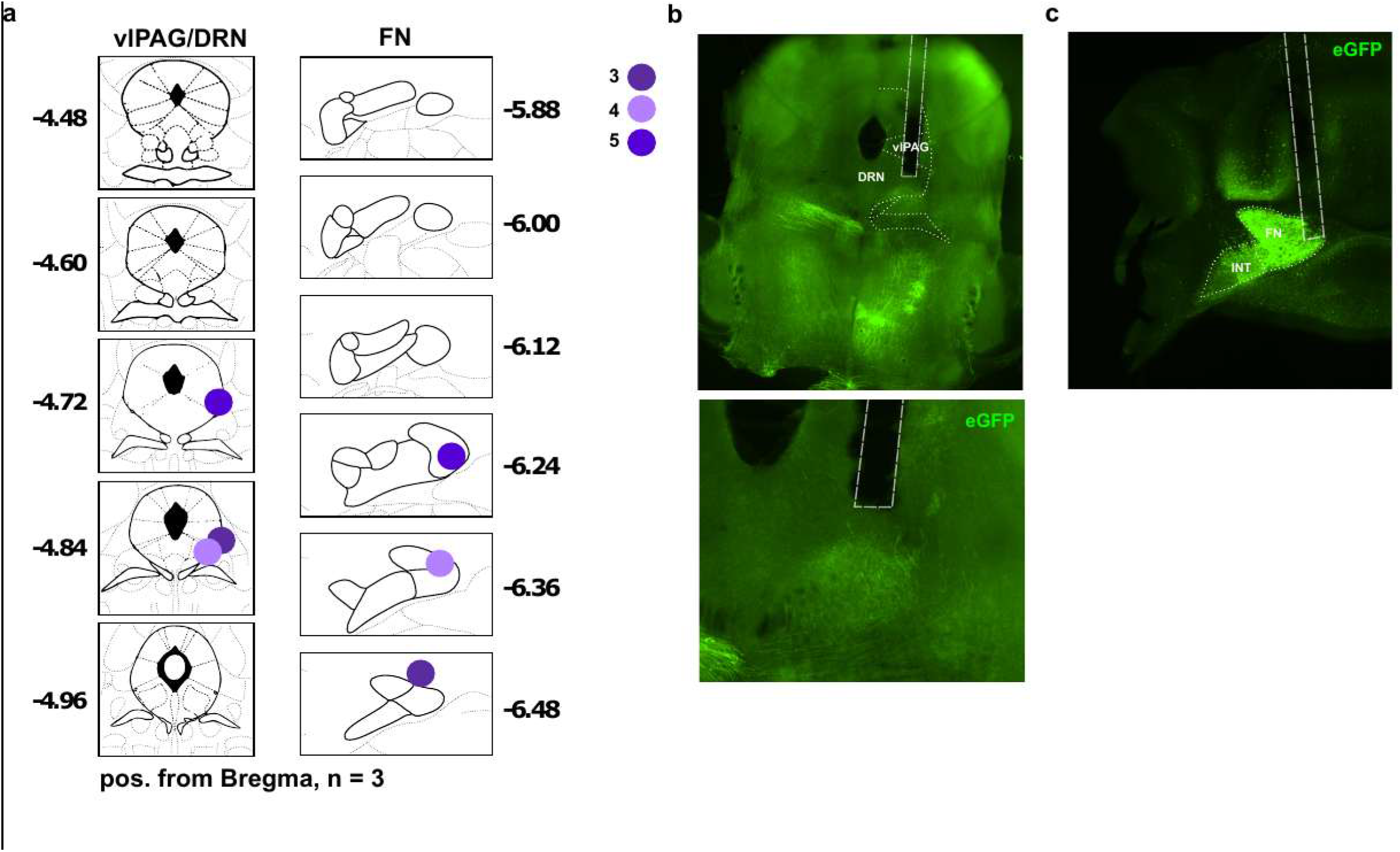
Excitatory response pattern in FN units stimulated by vlPAG/DRN to FN projecting neurons. (a) Optic fiber insertion sites in the vlPAG/DRN and electrode insertion sites in the FN. (b) Example of eGFP expression in the vlPAG in proximity to optic fiber, zoom in below. (c) Example of eGFP expression around electrode in FN.

### Spatial proximity of vlPAG/DRN to FN and FN to vlPAG/DRN projection neurons suggests reciprocal connections between cerebellum and midbrain

We then aimed to characterize the histological relationship between vlPAG/DRN to FN projection neurons and FN to vlPAG/DRN projection neurons to identify anatomical correlates of a reciprocal neural circuit between vlPAG/DRN and FN. Having confirmed the presence of vlPAG/DRN to FN projection neurons, we next examined whether these neurons are anatomically associated with the previously described FN to vlPAG/DRN monosynaptic projections that are functionally important for fear learning^23^.

To assess potential reciprocal connectivity, we designed a bidirectional transsynaptic viral assay to induce eGFP expression in vlPAG/DRN to FN projection neurons following transsynaptic transfer of Cre recombinase from FN neurons. Specifically, AAV1-pCAG-FLEX-eGFP-WPRE was injected into the vlPAG/DRN and AAV1-mCherry-cre in the FN (Fig. 6 a). Because both vectors possess transsynaptic properties, eGFP expression in vlPAG/DRN neurons should occur when Cre recombinase is transferred across synapses from FN neurons. These experiments may produce FN neurons expressing both mCherry and eGFP (appearing as yellow in merged images). Such co-expression indicates that these FN neurons receive inputs from the vlPAG/DRN neurons (Fig. 6 b left). Conversely, vlPAG/DRN neurons projecting to the FN and receiving transsynaptic Cre from FN neurons would express eGFP, resulting in eGFP-labeled fibers in the FN (Fig.6 b middle). Finally, the apposition of eGFP labeled fibers to eGFP/mCherry co expressing somata in the FN may provide anatomical evidence for a closed-loop connection between these FN neurons and vlPAG/DRN neurons (Fig.6 b right).

**Figure 6.**
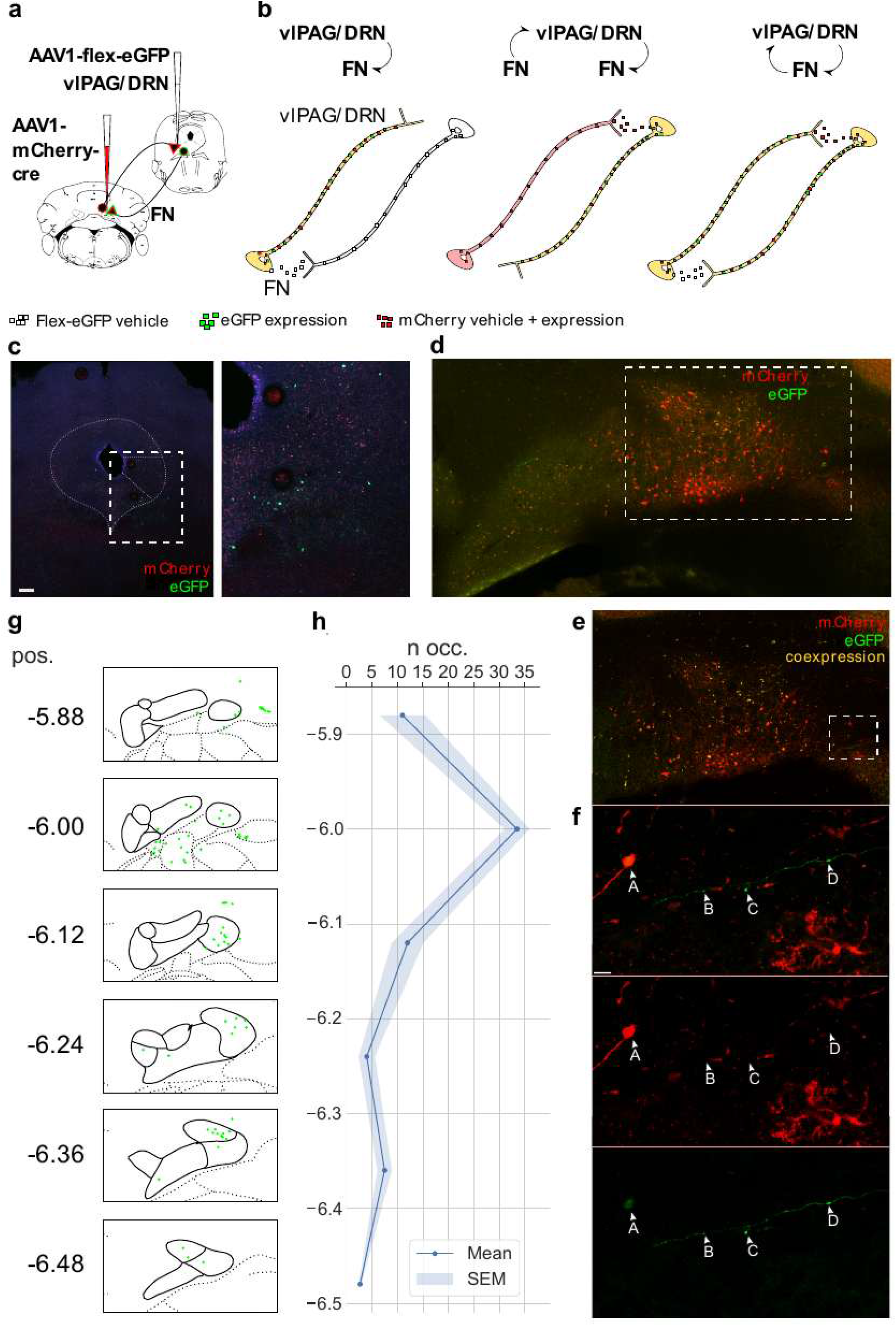
Neurons projecting from vlPAG/DRN to FN are in spatial proximity to FN to vlPAG neurons, revealing reciprocal projections between cerebellum and midbrain. (a) Schematic of the viral tracing strategy: injection of AAV1-CAG-flex-eGFP (1 × 10¹³ vg mL⁻¹) into the vlPAG/DRN and AAV1-hSyn-mCherry-Cre (1 × 10¹³ vg mL⁻¹) into the FN. (b) Schematics illustrating the resulting expression patterns in FN and vlPAG/DRN neurons. (c) Expression of eGFP and mCherry in the midbrain, with a zoom-in of the vlPAG/DRN region within a dashed rectangle. (d) Expression of eGFP and mCherry, the FN is highlighted by a dashed rectangle. (e) Magnified view of the region outlined in (D), showing mCherry, eGFP, and their co-expression. (f) Higher-magnification view of the dashed region in (E): arrows indicate eGFP- expressing structures (A, ∼10 µm; B, ∼1.1 µm; C, ∼2.3 µm; D, ∼2.8 µm). Middle, mCherry expression; top, eGFP/mCherry overlay. Scale bar, 10 µm. g) Distribution and quantification (n= 4) of eGFP/ mCherry co-expressing structures in coronal sections of the cerebellar nuclei and surrounding regions ipsilateral to the FN injection site, shown for a representative animal. (h) Co-expression quantification for all animals in coronal sections of the contralateral cerebellar /DRN.

We observed co-expression of eGFP and mCherry in both the FN and vlPAG/DRN (Fig. 6 c-e), indicating that Cre was expressed in vlPAG neurons following transsynaptic infection from FN neurons, consistent with the existence of FN neurons projecting to this area. Reciprocally, co-expression was notably observed in large somata in the ventral part of the FN (Fig. 6 f-g), and also in the neighboring vestibular nuclei (Fig. 6 g). In this assay, co-expressing FN neurons receive projections from vlPAG/DRN neurons expressing the FLEX-eGFP vector.

Strikingly, we also observed eGFP labeled fibers in the FN. These fibers may originate from the vlPAG/DRN neurons that received transsynaptic Cre from FN neurons projecting to the vlPAG/DRN. Alternatively, they may originate from another region in which vlPAG/DRN projection neurons and FN projection neurons converge onto a common neuron, allowing for transsynaptic transfer of the viral vector and subsequent labeling of neurons projecting back to the FN. In either case, this these observations are consistent with reciprocal connectivity between FN and vlPAG/DRN projecting neuronal populations. Moreover, we observed close apposition between such eGFP-labeled fibers and FN neurons expressing both eGFP and mCherry (Fig. 6 f, label A (soma) and labels B,C,D (fibers). This spatial proximity is consistent with the possibility that the FN neurons projecting to the vlPAG/DRN and vlPAG/DRN neurons projecting to the FN form a reciprocal circuit. However, the present anatomical approach does not by itself establish synaptic connectivity or distinguish between direct and indirect interactions.

## DISCUSSION

The neural circuitry underlying fear learning has been extensively described, yet the mechanisms underlying the computation of the prediction error, which drives the learning rate, remain largely unclear. Here, we provide evidence of functional coordination between the FN and vlPAG/DRN neuronal populations encoding the association between the CS+ and the aversive stimulus, reflecting associative fear learning. Moreover, we identified reciprocal projections from vlPAG/DRN to FN, in addition to the previously described FN to vlPAG/DRN projections, suggesting that bidirectional information flow may occur between the cerebellum and midbrain during fear learning and may contribute to predictive signaling. In parallel, both FN and vlPAG/DRN adapted their activity patterns to threat-related stimuli over the course of fear conditioning.

### Cerebellar inputs shape associative learning via predictive signaling

The cerebellum is increasingly recognized as an active contributor to emotional processing ^40,41^, including the formation and modulation of negative emotional memories^42^. More recently, the projections of the cerebellum to the vlPAG/DRN border region have been shown to modulate fear learning and extinction^23,25^. Analysis of the local field potentials evoked by the sound has been interpreted as consistent with an information flow from the FN to the vlPAG, which is inversely correlated with the degree of fear expression^43^. Consistent with these findings, we observed parallel development of CS+ evoked responses in the FN and vlPAG/DRN following fear conditioning; likely reflecting predictive signals associated with the aversive outcome. Furthermore, crosscorrelograms analysis revealed that, in a subset of neuron pairs, FN activity preceded vlPAG/DRN discharge; suggesting a temporal organization compatible with information transfer from the cerebellum to the midbrain. The origin of the CS+ response in the FN is unlikely to arise from the climbing fiber input from the inferior olive, as described in reward conditioning ^44,45^. Indeed, such recruitment typically involves complex spikes in Purkinje cells, resulting in transient inhibition of cerebellar nuclei activity, which was not observed in our recordings. While a transient reduction in Purkinje cell firing cannot be excluded, our data do not support a CS+ response driven exclusively by climbing fiber–driven modulation.

### Direct projections from the vlPAG/DRN to the FN

We identified direct anatomical projections from the vlPAG/DRN border region to FN neurons. As previously suggested^8,13^, this area may operate as an integrated hub recruited during both fear expression and fear learning. This observation is consistent with the established roles of the vlPAG in aversive associative learning^7,8^ and of the DRN in fear related processes^18,19^. Serotonergic neurons in the vlPAG/DRN increase their responsiveness to threat during fear conditioning^17–19^ and can promote behavioral responses appropriate to threat imminence based on their activity patterns^46^. Here we described a projection from the vlPAG/DRN to the FN. Such projection has been previously reported in the cat^37^ and its presence in the rodent brain indicates evolutionary conservation of this pathway. We further observed short latency FN responses following stimulation of the vlPAG/DRN projection neurons supporting a functional influence of this pathway. Interestingly, a polysynaptic pathway from the vlPAG to the cerebellar cortex via the inferior olive has also been described ^47^. However, this pathway has been associated primarily with fear expression rather than fear learning. Anatomically, these FN-projecting vlPAG/DRN neurons identified here appear to be located more ventral and lateral than the vlPAG/DRN neurons projecting to the central amygdala (CeA), as described by Groessl et al.^8^. Although their location resembles serotonergic FN projecting neurons reported by Kitzman in 1994^37^, we did not observe serotonin and dopamine reporter labeling in our FN-projecting vlPAG/DRN neurons suggesting that these cells may represent a non-monoaminergic population. We propose that these excitatory projections may be primarily glutamatergic, consistent with the predominance of glutamatergic neurons in the vlPAG^23,27^, while potentially being modulated by neighboring monoaminergic populations^27^. The vlPAG also interacts functionally with the cerebellar cortex during fear conditioning^47^, and several midbrain structures directly project to the cerebellar cortex^48^.

### Representation of predictive signal and fear related behavior in the vlPAG/DRN

Several neural populations within the vlPAG/DRN contribute to aversive behavior^15,27,46,47,49,50^, while others have been implicated in predictive signaling^7,8,20^. Dopaminergic neurons in the vlPAG/DRN have been reported to encode associative strength between CS and US, with their activity shifting toward the CS following conditioning ^8,11^. Neurons exhibiting predictive signals have also been described as dopaminergic^7^, which is consistent with the waveform characteristics observed in our PS population. These observations are consistent with the emergence of vlPAG/DRN PS units following fear conditioning. However, in our recordings, vlPAG/DRN PS neurons did not initially exhibit a strong response to the US, which is not consistent with a prediction error signal. We therefore hypothesize that the neurons identified here are not necessarily dopaminergic despite sharing some anatomical and electrophysiological characteristics with dopaminergic neurons. Their location within the vlPAG/DRN also places them in close proximity to serotoninergic populations raising the possibility that they may belong to, or be modulated by, serotoninergic circuits. Still, the absence of a strong initial response to the aversive US in PS neurons suggests that these neurons may not primarily encode the immediate aversive response itself. Instead, their activity may be more closely related to the learned expectation of an aversive event or to motivational processes associated with predictive signaling. Further characterization of their molecular and neurochemical identity will be necessary to distinguish between these possibilities.

### Cerebellar control over fear related behavior

Beyond predictive signaling, vlPAG/DRN neurons may encode a broader threat/fear state involving pain modulation^16,51^, anxiety related processes mediated by GABAergic interneurons and neuropeptide Y ^51^ and autonomic responses ^52^. We hypothesize that some of these functions may represented by the non-PS (nPS) population. Interestingly, we observed stable interactions between FN PS neurons and vlPAG/DRN nPS neurons, suggesting that predictive signals originating in the FN may influence vlPAG/DRN populations that do not themselves exhibit a predictive response. This interpretation is consistent with findings that the cerebellum can bidirectionally modulate fear learning^25,53^, and suggests that this influence may involve an FN population representing predictive signal. Thus, cerebellar predictive signaling may not only participate in the anticipation of aversive events but may also interact with broader threat-related neuronal populations in the vlPAG/DRN.

### Identity of neurons participating in the network exchange

Several neuronal populations are likely to contribute to the midbrain-cerebellar network described here. Studies in cats have identified vlPAG/DRN to FN projections as serotonergic^37^, while direct projections from the DRN to the cerebellar cortex have also been reported^54^. In our recordings, interactions involving FN neurons exhibited both excitatory and inhibitory temporal dynamics, with a predominance of patterns consistent with inhibitory coordination. Several mechanisms could contribute to this organization. First, direct projections from midbrain regions to the cerebellar cortex may influence FN activity indirectly to local cerebellar circuitry. Second, direct vlPAG/DRN projections to the FN may simultaneously influence FN neurons. The excitatory responses observed following optogenetic stimulation of vlPAG/DRN to FN projection neurons are consistent with the clear excitatory influence of this pathway. However, this alone does not explain the inhibitory like dynamics observed in the network-level cross-correlation analysis. One possibility is that vlPAG/DRN projections may engage inhibitory interneurons in the FN^55^. However, this mechanism would need to be reconciled with the net excitatory responses observed during optogenetic stimulation. Alternatively, the inhibitory like cross-correlation patterns may arise from indirect pathways, convergent inputs, or coordinated activity within broader neuronal network rather that from direct inhibitory synaptic transmission between recorded neuronal populations.

In the network dynamics observed during fear conditioning, FN activity also appeared to precede and coordinate with vlPAG/DRN activity. This observation is consistent with previous descriptions of FN to vlPAG/DRN projections that synapse on glutamatergic and dopaminergic neurons in the vlPAG/DRN^27,53^. Together, these findings support the existence of reciprocal communication between cerebellar and midbrain populations during fear learning.

### Circuit dynamics between vlPAG/DRN and FN may support mode switching between baseline and fear states

FN and vlPAG/DRN are reciprocally connected, as supported by our anatomical and functional experiments. This organization provides a substrate for reciprocal interactions that could dynamically influence behavior and learning. We observed that, during fear conditioning, interactions involving PS neurons become more prominent relative to the baseline behavioral state. Interestingly, the transition from baseline to fear conditioning appeared to involve broader interactions driven by FN PS neurons more than by the vlPAG/DRN PS neurons, but also with vlPAG/DRN populations that did not exhibit a predictive response. In contrast, interactions involving vlPAG/DRN PS neurons remained more closely associated with PS populations in the FN during both baseline and fear conditioning conditions. These observations suggests that fear conditioning may increase the functional relevance of predictive information within the FN - vlPAG/DRN network. In particular, FN neurons encoding predictive signals may influence a broader group of vlPAG/DRN populations, including neurons that do not themselves exhibit predictive responses. Such interactions could provide a mechanism through which cerebellar predictions are integrated with broader threat-related processes in the midbrain.

Although the anatomical basis for communication between FN and vlPAG/DRN is supported by direct projections in both directions, how these pathways are integrated to generate the functional interactions observed here remains unclear. The circuit is likely to involve multiple synaptic pathways and parallel projections. In addition, neuromodulatory influences, including dopaminergic and serotonergic signaling within the vlPAG/DRN, may contribute to the integration of fear learning and threat related information^18,50,56,57^.

### Further deciphering the network between FN and vlPAG/DRN

To determine whether and how this network contributes to the transition between baseline behavior and responsiveness to stimuli predicting aversive events, future studies should characterize the molecular and neurochemical identity of vlPAG/DRN neurons projecting to the FN and directly examine their contribution to fear learning and behavior. This would help determine the nature of the information communicated from the midbrain to the cerebellum in an aversive context, and distinguish the contribution of direct midbrain projections to the cerebellar nuclei from that of projections to the cerebellar cortex. For example, projections from the locus coeruleus to the cerebellar vermis^59^ produce patterns of noradrenaline release resembling the increase in FN firing-rate that we observed during fear conditioning. This raises the possibility that noradrenergic signaling may contribute to the modulation or gating of cerebellar activity during aversive learning^59^.

### Implications for maladaptive threat processing

In summary, we propose that the cerebellum contributes to associative fear learning by providing predictive signals related to the expectation of aversive events, while its interactions with midbrain structures such as the vlPAG/DRN are recruited during exposure to aversive stimuli. Reciprocal communication between these regions provides a potential substrate for cerebellar - midbrain interactions that are distinct from the inferior olive pathway and may contribute to adaptive fear learning.

This raises the possibility that the vlPAG/DRN–FN circuit could be relevant to disorders involving maladaptive threat processing. Future studies manipulating specific parts of the midbrain-cerebellar circuit will be required to determine whether altering these interactions can modify the anticipation of aversive events and associated behavioral responses, and whether such mechanisms could ultimately have therapeutic relevance.

## MATERIAL AND METHODS

### Animals

Adult male C57BL/6N mice (Charles River Laboratories, France) were used for all experiments. Animals were housed in groups of four per cage under controlled environmental conditions (21–22 °C, 40% humidity, 12 h light/dark cycle) with food and water available ad libitum. One week before fear conditioning, animals were individually housed to facilitate handling and behavioral monitoring. Only adult mice aged 8–12 weeks were included in the study. Male and female mice can both be used for fear conditioning studies; however, we used only male mice to minimize potential variability associated with the estrous cycle over the course of our behavioural paradigm. Fluctuations in ovarian hormone levels across the estrous cycle can modulate neural excitability and behavior, particularly in circuits involved in affective and stress-related processing. Estrous stage-dependent changes in neuronal excitability and anxiety-related behavior have been reported in multiple brain regions, including limbic and midbrain circuits, with measurable effects on physiological and behavioral readouts across the cycle ^60^. The design choice was therefore intended to reduce a potential source of variability within the experimental cohort rather than to imply sex-specific differences in the suitability of the fear-conditioning paradigm. Animal care and experimental procedures were conducted in accordance with the European Community Council Directives and applicable institutional and national regulations (authorization number APAFIS#1334-2015070818367911 v3 and APAFIS #29793-202102121752192).

### Surgeries

Surgeries for tracing and electrophysiological experiments were performed in 7–8 weeks-old mice. Animals were placed in a stereotaxic frame (Kopf Instruments) following buprenorphine administration and maintained under isoflurane anesthesia (2%) throughout the whole procedure. Local analgesia was additionally provided by subcutaneously administration of 0.02% lidocaine above the skull. The body temperature was continuously monitored and maintained at 37 °C using a heating pad and a rectal thermometer. For viral injections and electrode implantations, stereotaxic coordinates (in mm) for the FN (AP: −6.43 to 6.49, ML: ±0.77, DV: −2.14 to 2.18) and vlPAG/DRN (AP: −4.45 to 4.49, ML: ±0.55, DV: −2.45 to 2.49) were measured relative to bregma, with the dorsoventral coordinate measured from the brain surface. For viral delivery, a quartz capillary was lowered to a position immediately above the target structure, and 150–250 nl of virus solution was infused. Quartz capillaries (World precision instruments, QTZ,1.0x.50mmx7.5cm, FILMT) were left in place for 5 minutes after the injection before being slowly withdrawn.

### Histology

At the conclusion of the experiments, mice were deeply anesthetized with a mixture of zoletil/xylazine (50 and 10 mg/kg, i.p., respectively) and rapidly transcardially perfused with ice-cold 4% paraformaldehyde in phosphate buffered SAL (PBS). Brains were dissected and post-fixed overnight at 4 °C in 4% paraformaldehyde, then stored in PBS at 4 °C until sectioning. Brains were sectioned throughout their entire extent at 90 µm using a vibratome (Leica VT 1000 S). Sections were mounted on gelatin-coated slides, dried, and cover slipped with Mowiol (Sigma). Sections were imaged using a confocal microscope (Leica TCS Sp8), and images were processed and analyzed using Image J. Electrode placement and viral injection sites were assessed histologically for all the experiments at the end of the experimental procedures. Animals with no detectable viral expression, or incorrect electrode placement were excluded from the analysis.

### Fear conditioning and recall

For electrophysiological recordings, the following behavioral paradigm was used in freely moving animals. 3 days before to the conditioning paradigm, all animals underwent an assessment of anxiety related behavior. On the first experimental day, animals underwent a CS-habituation session in the morning and fear conditioning in the afternoon. On the second experimental day, animals underwent a recall session.

The CS habituation session consisted in 5 minutes of acclimatization followed by 4 CS- and 4 CS+ presentations, with an inter-trial-interval (ITI) of 20-40 s, and 5 minutes of rest in the experimental chamber. The fear conditioning session consisted of 5 minutes of acclimatization followed by 5 CS+ presentations co-terminating with a foot shock and 5 CS- presentations, delivered in a pseudo-random order with an ITI of 60-120 s and 5 minutes of rest in the experimental chamber. The session always began with a CS+ presentation co-terminating with the foot shock. Recall sessions consisted of 5 minutes of acclimatization followed by 4 CS- and 12 CS+ presentations without US delivery, with an ITI of 20-40 s, and 5 minutes of rest in the experimental chamber.

Two distinct types of CS that were used: 30 s white noise at approximately 65 dB and a 30 s pure tone at 7000Hz and 70 dB. Half of the animals received white noise as the CS+, whereas the other half received the pure tone as the CS+. The US consisted of a 0.5 s, 0.4 mA square wave electrical foot shock delivered through an electrified floor grid at 29.5 s after CS+ onset, such as the shock co-terminated with the CS+. Stimulus presentation was controlled using the Ethovision XT 14 (Noldus).

Fear conditioning and recall were conducted in two distinct contexts (A and B), both located inside of a sound attenuating box (Ugo Basile). CS habituation and fear conditioning were conducted in context A. consisting of a 17 × 17 × 25 cm square chamber with black and white-checkered walls and a grid floor, scented with peppermint-soap. The chamber was cleaned with 70% ethanol between animals. Recall was conducted in context B, consisting of a 17-cm diameter yellow plexiglass cylinder with a grey floor and vanilla odor. This champer was cleaned with 1% acetic acid between animals.

### Anatomical tracing

For exploratory neuroanatomical tracing, a group of mice (n=2) received an injection of 150 nL of pENN.AAV.hSyn.TurboRFP.WPRE.RBG (AAV1, Addgene, 105552-AAV1) in the vlPAG/DRN. For double virus neuroanatomical tracing, an injection of 150-250 nL of pAAV-Ef1a-mCherry-IRES-Cre (AAV Retrograde, Addgene, 55632-AAVrg) into the FN was combined with an injection of 150-250 nL of AAV-pCAG-FLEX-eGFP-WPRE (AAV1, Addgene, 51502-AAV1) into the vlPAG/DRN in male C57Bl6N mice (n = 3). For double virus circuit trans-synaptic labeling, a group of mice received an injection of 150-250 nL of AAV-pCAG-FLEX-eGFP-WPRE (AAV1, Addgene, 51502-AAV1) into the vlPAG/DRN (concentration 10^13), together with an injection of 150-250 nL of AAV1-hSyn-mCherry-P2A-Cre-WPRE (AAV1, Addgene, 107312-AAV1) into the FN.

Mice were deeply anesthetized with zoletil (50 mg kg−1) and xylazine (10 mg kg−1 i.p.), and transcardially perfused with formalin (Sigma) 3–4 weeks after surgery to assess the viral expression. Brains were dissected and post-fixed overnight in formalin at 4 °C and then transferred to PBS until sectioning. Coronal brain sections (50 or 90 μm) were obtained using a vibratome (Leica VT 1000 S), then dried and mounted with Mowiol (Sigma). Sections were analyzed and imaged using a confocal microscope (Leica TCS Sp8).

### Immunostaining

For the immunostaining, 50 µm thick sections containing the vlPAG/DRN were washed with PBS-0.3% Triton X-100. Non-specific binding sites were blocked by incubating the sections for 2 h in PBS containing 3% normal donkey serum (NDS). Sections were then incubated overnight at 4°C in PBS containing 0.3 % Triton X-100 and 1.5% NDS with either a polyclonal rabbit anti-TPH2 antibody (1:400, ThermoFisher PA1778) or a polyclonal guinea pig anti-Tyrosine hydroxylase antibody (1:300, Synaptic systems, 213104). Following primary antibody incubation, sections were rinsed and incubated for 2 h at room temperature with the appropriate secondary antibody: goat anti-rabbit IgG conjugated to Alexa Fluor 594 (1:300, ThermoFisher, A11012) or CyTM3-conjugated donkey anti-guinea pig IgG (1:300, Jackson Immuno Research, 706-165-148). Sections were then mounted with Mowiol and imaged using a confocal microscope (Leica TCS Sp8).

### Optogenetics and electrophysiological recordings

For optical manipulation combined with electrophysiological recordings, mice were injected bilaterally with 200 nL of ssAAV-retro/2-hSyn1-hChR2(H134R)_EYFP-WPRE-hGHp(A) (Zuerich vial facility, v221-retro) in the FN. For optical stimulation, mice were implanted unilaterally with an optic fiber (0.22 numerical aperture, Thorlabs) housed in a stainless steel ferrule (ThorLabs) positioned just dorsal to the vlPAG/DRN. Ferrules were secured to the skull with adhesive, and a protective head cap was constructed using dental cement. Because the vlPAG/DRN preferentially projects to the contra-lateral FN, neuronal activity was recorded in the contralateral FN while the vlPAG/DRN was unilaterally stimulated. To record neuronal activity in the FN, we used bundle of electrodes consisting of nickel/Chrome wire (0.004 inches diameter, Coating 1/4 Hard PAC) folded and twisted into bundles of six electrodes. Pairs of electrode bundles were inserted into metal cannulas (stainless steel, 29 G, 8 mm length, Phymep), to protect the electrodes and ensure accurate placement into the brain. Cannulas and electrode bundles were attached to an electrode interface board (EIB-16; Neuralynx, Bozeman) using Loctite universal glue, in a configuration allowing recordings from the FN. The microwires of each bundle were connected to the EIB using gold pins (Neuralynx). The EIB and all connections were secured in place with dental cement (Super Bond). Electrode impedance was measured at electrodes were gold-plated using a cyanure-free gold solution (Sifco) to achieve an impedance of 200–500 kΩ. During surgery, the electrode bundles were inserted into the brain and the ground was positioned over the cerebellum, between the dura and the skull. Viral injections, optical fiber implantations, and electrodes implantation were performed during the same surgery. Optical stimulation and electrophysiological recordings were performed 3 weeks after surgery to allow sufficient time for ChR2 expression. At the end of the experiments, optic fiber placement, viral expression and electrode insertion sites were histologically verified.

For the optical stimulation intensity ramp, electrophysiological recordings in the FN were performed in freely moving mice placed in an open field arena. Recordings were performed using a 16 channel acquisition system (sampling rate 25 kHz; Tucker-Davis Technology System 3, Tucker-Davis Technologies (TDT)), as previously described^61,62^. Mice received optical stimulation at 0.5 Hz with a pulse duration of 100 ms. Light intensities of 39.8, 49, 67.8 and 83.3 mW/mm2 at the fiber tip were used. Signals were acquired using a headstage and amplifier from TDT (RZ2, RV2, Tucker-Davis Technologies). Spike sorting was performed with Mountainsort4^63^ (Version: 1.0.4, spike sorting algorithm) in spike interface^64^ (Version: 0.97.1, python toolkit).

### Electrode implantation and chronic electrophysiological recordings

Extracellular recordings were performed using standard 16 channel electrode interface boards (EIB-16; Neuralynx), to which 2 cannulas were attached, one targeting each recorded region, the vlPAG/DRN and FN. Electrode bundles consisted of nickel chrome wires (16μm diameter, Coating ¼ Hard PAC, KANTHAL Precision Technology) twisted in bundles of 6 wires and gold-plated to achieve an impedance of 200–500 kΩ (cyanure-free gold solution, Sifco). One or two bundles were inserted into each cannula (stainless steel, 30 Gauge, Phymep).

The recording electrodes were progressively lowered until they reached the targeted brain structures, and the electrode interface boards were secured to the skull using dental adhesive resin cement (Super-Bond C&B, Parkell). Miniature stainless-steel screws were implanted on the left parieto-occipital suture, serving as electrical reference and ground. The skin was sutured, and mice were allowed to recover in their home cage for at least 2 weeks. Recordings were performed using a wireless head stage equipped with a battery and connected to a Multi-Channel System W2100 system (sampling frequency: 25000 Hz) using MCS recording software (v1.5.6). Mice were habituated to the head-stage and its weight for 10 minutes per day starting one week before the experiment.

### Behavioral analysis

Immobility was quantified from video recordings acquired using a Basler acA 1300-60gm camera mounted on the same Ugo Basile conditioning behavioral setup and controlled using EthoVision XT 14 software. During electrophysiological recordings, immobility levels during fear conditioning were detected using the video analysis tools in EthoVision XT,. Periods of immobility lasting longer than 2 s during CS presentation were classified as immobility. For contextual immobility levels during habituation and fear conditioning on experimental day 1, the percentage of time spent immobile during the first 5 minutes preceding CS presentation were classified as immobility.

### Statistical analysis

#### Behavioral data statistical analysis

To assess group differences between experimental and control animals, an ANOVA was performed to compare immobility percentages during the CS+ presentations during behavior. Analysis was conducted in python using SciPy (Version: 0.13.2) statsmodels (Version: 0.13.2) packages.

#### Electrophysiological signal analysis

Electrophysiological recordings were band-pass filtered to retain frequencies between 300 and 8000 Hz and subsequently sorted into single units. Spike sorting was performed using Mountainsort4^63^, with standard settings and a detection threshold of 4 (Version: 1.0.4, spike sorting algorithm) in spike interface^64^ (Version: 0.97.1, python toolkit). Single units identified by Mountainsort were then filtered according to their firing rate (above 1 and below 150 Hz), waveform characteristics and autocorrelogram shape. Units were retained if their waveform exhibited a trough and a peak separated by a maximum of 2 ms, a maximum halfwidth of 0.6 ms, and a maximum peak/trough ratio of 0.85. Units showing visible noise contamination in their peri-stimulus time histograms (PSTHs) were excluded for subsequent analysis (see Supplementary information). For waveform shape clustering, waveform halfwidth, amplitude ratio and recovery time were computed for each unit. The amplitude ratio was calculated as trough-peak/(trough+peak), while recovery time was defined as the interval from the trough to recovery to baseline. Principal component analysis (PCA) was performed on these parameters, and the optimal number of clusters was determined using a silhouette test. K-means clustering was then performed using scikit-learn Python package (version 1.7.2.)

#### Optogenetic stimulation coupled with electrophysiological signal analysis

To analyze neuronal responses to the optogenetic stimulation, peri-stimulus time histograms (PSTHs, 1 ms bins) were constructed for each stimulation event. For each stimulation, a baseline window was defined as the 100 ms preceding stimulation onset, and the response window was defined as the 100 ms following stimulation onset. To identify stimulation responsive units, mean firing rate (FR) during the baseline and response windows was compared for each unit using a paired t-test (SciPy version: 0.13.2). To visualize local field potential (LFP) deflections associated with optogenetic stimulation, recordings were butterwoth-band-pass filtered between 0.5 and 300Hz. PSTHs were then constructed around the stimulation onset for each electrode channel and plotted as the mean LFP deflection +/- SEM.

#### Single structure single unit electrophysiological analysis

Firing rates were computed from spike-sorted single unit recordings as described above. For analyses performed separately in the vlPAG/DRN and FN analysis windows were centered on CS+ onset, CS- onset or US delivery. For CS onset, the first 5 s following CS onset were analyzed. For US-evoked activity, a 5 s window following US delivery was used, excluding the US itself to avoid stimulation artifacts. These analysis windows are referred to as reporter time windows (RTWs). A 20 s period preceding CS onset were used as the baseline. To visualize population activity during CS and US presentations, peri stimulus time histograms (PSTHs, 1 s bin size), were computed and normalized by subtracting the baseline firing rate. The ‘naïve’ condition was defined as the firing rate of each unit during the first CS+ or CS- presentation or the first US presentation. The ‘trained’ condition was defined as the mean firing rate across the three last presentations of the corresponding stimulus (the 3^rd^,4^th^ and 5^th^ presentations).

For naïve versus trained comparisons, the mean firing rate within each reporter time window (RTW) was calculated for each unit after subtraction of the baseline firing rate. Naïve and trained responses were then compared using Wilcoxon-rank test (R stats package version 4.1.2). The change in firing rate (Delta FR) was computed for each unit by subtracting the naïve response from the trained response. To classify units as PS and non-PS units, k-means clustering was performed on the Delta FR values calculated within the 2 to 6 seconds window following CS onset. Differences in responses between PS and non-PS units during CS+ or CS- presentations or following US delivery were assessed using the mean FR within the corresponding reporter time windows (RTW) and the Wilcoxon rank test. The analysis were performed in R (version 4.1.2 (2021-11-01)) using Rstudio (2023.12.1.402).

#### Connectivity analysis

Cross-correlograms were computed between simultaneously recorded units from different structures within the same animal according to their cluster affiliation. Correlograms were calculated for PS – PS pairs, non-PS – non-PS pairs and PS – non-PS pairs. Spike-trains were analysed during 2 time windows: baseline, define as the 5 minutes preceding the first CS presentation and fear conditioning, emcompassing the period from the first to last CS presentation. Cross-correlograms were computed using 3 ms bins over a window of 300 ms. Cross-correlograms were then summarized in a matrices according to neuronal pairs type and behavioral condition. Asymmetry was quantified by comparing the number of spikes in the 27 ms preceding and following the reference unit discharge. The statistical significance of asymmetry was assessed using a Wilcoxon signed-rank test against zero. Significant pairs were classified according to the direction of asymmetry: positive asymmetry was interpreted as the vlPAG/DRN unit preferentially firing after the FN reference unit, and negative asymmetry as the vlPAG/DRN unit preferentially firing before the FN reference unit.

For the analysis of changes in asymmetry across behavioral conditions, mean asymmetry values for each neuronal pair subgroup were compared between baseline and fear conditioning using ANOVA over the 15-ms period following reference unit discharge. For comparisons of asymmetry between neuronal subgroups within the same time window, group-averaged cross-correlograms were visualized over a 20-ms window before and after reference unit discharge using 5-ms bins. Statistical comparisons were performed on the summed asymmetry values across the first three 3-ms bins (9 ms in total) following reference unit discharge using the Wilcoxon rank-sum test. Statistical analyses were performed in R using the stats package (version 4.1.2).

#### Sample sizes and statistical tests

For the electrophysiological analysis, sample sizes were determined by the number of neurons reliably recorded from the experimental animals and were comparable to those used in previous in vivo electrophysiological studies^24,53^. Given the relatively small sample sizes and the paired nature of the observations, differences between time points were assessed using the non-parametric Wilcoxon signed-rank test, which does not require the assumption of normally distributed data. For the asymmetry analyses, statistical significance was similarly assessed using a Wilcoxon signed-rank test against zero. For group comparisons, repeated-measures ANOVAs were used to assess the effect of condition over time. The same animals were repeatedly assessed across different conditions and/or time points, allowing within-subject changes and condition-by-time interactions to be evaluated while accounting for the paired nature of the observations. For anatomical studies, sample sized were determined based on the number of successful tracings and staining experiments and were comparable to those reported in previous studies^24,53^.

## DATA AVAILABILITY

Our data will be made available in an online repository upon acceptance of the manuscript.

## AUTHOR CONTRIBUTIONS

D.P. and C.L. acquired the funding and supervised the project; J. D. U. D. performed experiments, data curation and analysis. All authors contributed to the design, interpreted results, participated to the writing and approved the final manuscript.

## ACKNOWLEDGEMENTS AND FUNDING INFORMATION

This work was supported by Agence Nationale de Recherche to D.P (ANR-21-CE37-0025 CerebellEMO, ANR-24-CE37-3944 Ce-Multi-TimeS) to C.L. (ANR-21-CE16-0036 IntTempComp, ANR-21-CE16-0017 PomPom), by Fondation pour la Recherche Médicale FRM EQU-202103012770 and EQU-202603051384, and by the CEN European Union’s Horizon 2020 Research and Innovation Programme under the Marie Skłodowska-Curie grant agreement No. 956414 and by the Institut National de la Santé et de la Recherche Médicale (France) to D.P. and C.L.; to J.D.U.D. by the FRM FDT202404018688. This work used the IBENS imaging facility (IMACHEMIBiSA), member of the French National Research Infrastructure France-BioImaging (ANR-10-INBS-04), which received support from the "Fédération pour la Recherche sur le Cerveau – Rotary International France" (2011) and from the program « Investissements d’Avenir » ANR-10-LABX-54 Memolife.

## CONTACT AND COMPETING INFORMATION

The authors declare that they have no competing interests.

## Notes

### Competing Interest Statement

The authors have declared no competing interest.

## BIBLIOGRAPHY

1. Blanchard, D.C., and Blanchard, R.J. (1988). Ethoexperimental Approaches to the Biology of Emotion. Annu. Rev. Psychol. 39, 43–68. 10.1146/annurev.ps.39.020188.000355.

2. Fanselow, M.S. (1994). Neural organization of the defensive behavior system responsible for fear. Psychon. Bull. Rev. 1, 429–438. 10.3758/BF03210947.

3. Pavlov, I.P. (1927). Conditioned reflexes: an investigation of the physiological activity of the cerebral cortex (Oxford Univ. Press).

4. Pearce, J.M., and Hall, G. (1980). A model for Pavlovian learning: Variations in the effectiveness of conditioned but not of unconditioned stimuli. Psychol. Rev. 87, 532–552. 10.1037/0033-295X.87.6.532.

5. Rescorla, R., and Wagner, A. (1972). A theory of Pavlovian conditioning: The effectiveness of reinforcement and non-reinforcement. Classical Conditioning: Current Research and Theory.

6. Quirk, G.J., Repa, J.C., and LeDoux, J.E. (1995). Fear conditioning enhances short-latency auditory responses of lateral amygdala neurons: Parallel recordings in the freely behaving rat. Neuron 15, 1029–1039. 10.1016/0896-6273(95)90092-6.

7. Johansen, J.P., Tarpley, J.W., LeDoux, J.E., and Blair, H.T. (2010). Neural substrates for expectation-modulated fear learning in the amygdala and periaqueductal gray. Nat. Neurosci. 13, 979–986. 10.1038/nn.2594.

8. Groessl, F., Munsch, T., Meis, S., Griessner, J., Kaczanowska, J., Pliota, P., Kargl, D., Badurek, S., Kraitsy, K., Rassoulpour, A., et al. (2018). Dorsal tegmental dopamine neurons gate associative learning of fear. Nat. Neurosci. 21, 952–962. 10.1038/s41593-018-0174-5.

9. Apps, R., and Strata, P. (2015). Neuronal circuits for fear and anxiety — the missing link. Nat. Rev. Neurosci. 16, 642. 10.1038/nrn4028.

10. Tovote, P., Fadok, J.P., and Lüthi, A. (2015). Neuronal circuits for fear and anxiety. Nat. Rev. Neurosci. 16, 317–331. 10.1038/nrn3945.

11. Mcnally, G., Johansen, J., and Blair, H. (2011). Placing prediction into the fear circuit. Trends Neurosci. 34, 283–292. 10.1016/j.tins.2011.03.005.

12. Luo, M., Zhou, J., and Liu, Z. (2015). Reward processing by the dorsal raphe nucleus: 5-HT and beyond. Learn. Mem. 22, 452–460. 10.1101/lm.037317.114.

13. Sengupta, A., and Holmes, A. (2019). A discrete dorsal raphe to basal amygdala 5-HT circuit calibrates aversive memory. Neuron 103, 489–505.e7. 10.1016/j.neuron.2019.05.029.

14. Wright, K.M., and McDannald, M.A. (2019). Ventrolateral periaqueductal gray neurons prioritize threat probability over fear output. Elife 8. 10.7554/ELIFE.45013.

15. Wright, K.M., Jhou, T.C., Pimpinelli, D., and McDannald, M.A. (2019). Cue-inhibited ventrolateral periaqueductal gray neurons signal fear output and threat probability in male rats. Elife 8, e50054. 10.7554/eLife.50054.

16. Yan, C., and Liu, Z. (2024). The role of periaqueductal gray astrocytes in anxiety-like behavior induced by acute stress. Biochem. Biophys. Res. Commun. 720, 150073. 10.1016/J.BBRC.2024.150073.

17. Koutsikou, S., Crook, J.J., Earl, E. V, Leith, J.L., Watson, T.C., Lumb, B.M., and Apps, R. (2014). Neural substrates underlying fear-evoked freezing: the periaqueductal grey–cerebellar link. J. Physiol. 592, 2197–2213. 10.1113/jphysiol.2013.268714.

18. Yu, W., Pati, D., Pina, M.M., Schmidt, K.T., Boyt, K.M., Hunker, A.C., Zweifel, L.S., McElligott, Z.A., and Kash, T.L. (2021). Periaqueductal gray/dorsal raphe dopamine neurons contribute to sex differences in pain-related behaviors. Neuron 109, 1365–1380.e5. 10.1016/J.NEURON.2021.03.001/ASSET/4A5BB203-0327-40B2-9CF7-02922E77EAD3/MAIN.ASSETS/GR8.JPG.

19. Xie, L., Wu, H., Chen, Q., Xu, F., Li, H., Xu, Q., Jiao, C., Sun, L., Ullah, R., and Chen, X. (2022). Divergent modulation of pain and anxiety by GABAergic neurons in the ventrolateral periaqueductal gray and dorsal raphe. Neuropsychopharmacology 2022 48:10 48, 1509–1519. 10.1038/s41386-022-01520-0.

20. Ozawa, T., Ycu, E.A., Kumar, A., Yeh, L.-F., Ahmed, T., Koivumaa, J., and Johansen, J.P. (2017). A feedback neural circuit for calibrating aversive memory strength. Nat. Neurosci. 20, 90–97. 10.1038/nn.4439.

21. Ernst, T.M., Brol, A.E., Gratz, M., Ritter, C., Bingel, U., Schlamann, M., Maderwald, S., Quick, H.H., Merz, C.J., and Timmann, D. (2019). The cerebellum is involved in processing of predictions and prediction errors in a fear conditioning paradigm. Elife 8, e46831. 10.7554/eLife.46831.

22. Hoche, F., Guell, X., Vangel, M.G., Sherman, J.C., and Schmahmann, J.D. (2018). The cerebellar cognitive affective/Schmahmann syndrome scale. Brain 141, 248. 10.1093/brain/awx317.

23. Mark, M., Pakusch, J., Ernst, T.M., and Timmann, D. (2022). Cerebellum and Emotion Memory. In The Emotional Cerebellum Advances in Experimental Medicine and Biology., M. Adamaszek, M. Manto, and D. J. L. G. Schutter, eds. (Springer International Publishing), pp. 53–73. 10.1007/978-3-030-99550-8_5.

24. Stoodley, C.J. (2012). The cerebellum and cognition: evidence from functional imaging studies. Cerebellum 11, 352–365. 10.1007/s12311-011-0260-7.

25. Hoche, F., Guell, X., Vangel, M.G., Sherman, J.C., and Schmahmann, J.D. (2018). The cerebellar cognitive affective/Schmahmann syndrome scale. Brain 141, 248. 10.1093/brain/awx317.

26. Schmahmann, J.D., and Sherman, J.C. (1997). Cerebellar cognitive affective syndrome. Int. Rev. Neurobiol. 41, 433–440. 10.1016/s0074-7742(08)60363-3.

27. Schmahmann, J.D., and Sherman, J.C. (1998). The cerebellar cognitive affective syndrome. Brain 121 (Pt 4), 561–579. 10.1093/brain/121.4.561.

28. Frontera, J.L., Sala, R.W., Georgescu, I.A., Baba Aissa, H., d’Almeida, M.N., Popa, D., and Léna, C. (2023). The cerebellum regulates fear extinction through thalamo-prefrontal cortex interactions in male mice. Nat. Commun. 14, 1508. 10.1038/s41467-023-36943-w.

29. Frontera, J.L., Aissa, H.B., Sala, R.W., Mailhes-Hamon, C., Georgescu, I.A., Léna, C., and Popa, D. (2020). Bidirectional control of fear memories by cerebellar neurons projecting to the ventrolateral periaqueductal grey. Nat. Commun. 11, 1–17. 10.1038/s41467-020-18953-0.

30. Carta, I., Chen, C.H., Schott, A.L., Dorizan, S., and Khodakhah, K. (2019). Cerebellar modulation of the reward circuitry and social behavior. Science 363. 10.1126/science.aav0581.

31. Lawrenson, C., Paci, E., Pickford, J., Drake, R.A.R., Lumb, B.M., and Apps, R. (2022). Cerebellar modulation of memory encoding in the periaqueductal grey and fear behaviour. Elife 11, e76278. 10.7554/eLife.76278.

32. Koutsikou, S., Watson, T.C., Crook, J.J., Leith, J.L., Lawrenson, C.L., Apps, R., and Lumb, B.M. (2015). The Periaqueductal Gray Orchestrates Sensory and Motor Circuits at Multiple Levels of the Neuraxis. The Journal of Neuroscience 35, 14132–14147. 10.1523/JNEUROSCI.0261-15.2015.

33. Vaaga, C.E., Brown, S.T., and Raman, I.M. (2020). Cerebellar modulation of synaptic input to freezing-related neurons in the periaqueductal gray. Elife 9, e54302. 10.7554/eLife.54302.

34. Watson, T., Becker, N., Apps, R., and Jones, M. (2014). Back to front: cerebellar connections and interactions with the prefrontal cortex. Front. Syst. Neurosci. 8.

35. Salinas-Hernández, X.I., Vogel, P., Betz, S., Kalisch, R., Sigurdsson, T., and Duvarci, S. (2018). Dopamine neurons drive fear extinction learning by signaling the omission of expected aversive outcomes. Elife 7, e38818. 10.7554/eLife.38818.

36. Kitzman, P.H., and Bishop, G.A. (1994). The origin of serotoninergic afferents to the cat’s cerebellar nuclei. Journal of Comparative Neurology 340, 541–550. 10.1002/cne.903400407.

37. Jin, X., and Costa, R.M. (2010). Start/Stop Signals Emerge in Nigrostriatal Circuits during Sequence Learning. Nature 466, 457. 10.1038/NATURE09263.

38. Liu, X., Huang, H., Snutch, T.P., Cao, P., Wang, L., and Wang, F. (2022). The Superior Colliculus: Cell Types, Connectivity, and Behavior. Neuroscience Bulletin 2022 38:12 38, 1519–1540. 10.1007/S12264-022-00858-1.

39. Stoodley, C.J., and Schmahmann, J.D. (2009). Functional topography in the human cerebellum: a meta-analysis of neuroimaging studies. Neuroimage 44, 489–501. 10.1016/j.neuroimage.2008.08.039.

40. Guell, X., Gabrieli, J.D.E., and Schmahmann, J.D. (2018). Triple representation of language, working memory, social and emotion processing in the cerebellum: convergent evidence from task and seed-based resting-state fMRI analyses in a single large cohort. Neuroimage 172, 437–449. 10.1016/J.NEUROIMAGE.2018.01.082.

41. Damasio, A.R., Grabowski, T.J., Bechara, A., Damasio, H., Ponto, L.L., Parvizi, J., and Hichwa, R.D. (2000). Subcortical and cortical brain activity during the feeling of self-generated emotions. Nat. Neurosci. 3, 1049–1056. 10.1038/79871.

42. Paci, E., Lumb, B.M., Apps, R., Lawrenson, C.L., and Moran, R.J. (2023). Dynamic causal modeling reveals increased cerebellar-periaqueductal gray communication during fear extinction. Front. Syst. Neurosci. 17, 1148604. 10.3389/FNSYS.2023.1148604/BIBTEX.

43. Kostadinov, D., Beau, M., Blanco-Pozo, M., and Häusser, M. (2019). Predictive and reactive reward signals conveyed by climbing fiber inputs to cerebellar Purkinje cells. Nat. Neurosci. 22, 950–962. 10.1038/s41593-019-0381-8.

44. Jin, S., and Hull, C. (2025). Reward-driven cerebellar climbing fiber activity influences both neural and behavioral learning. Current Biology 35, 4383–4393.e4. 10.1016/j.cub.2025.07.064.

45. Stanley, A.T., Post, M.R., Lacefield, C., Sulzer, D., and Miniaci, M.C. (2023). Norepinephrine release in the cerebellum contributes to aversive learning. Nat. Commun. 14, 1–11. 10.1038/S41467-023-40548-8;TECHMETA.

46. Seo, C., Guru, A., Jin, M., Ito, B., Sleezer, B.J., Ho, Y.Y., Wang, E., Boada, C., Krupa, N.A., Kullakanda, D.S., et al. (2019). Intense Threat Switches Dorsal Raphe Serotonin Neurons to a Paradoxical Operational Mode. Science 363, 538. 10.1126/SCIENCE.AAU8722.

47. Zhang, H., Li, L., Zhang, X., Ru, G., and Zang, W. (2024). Role of the Dorsal Raphe Nucleus in Pain Processing. Brain Sciences 2024, Vol. 14, Page 982 14, 982. 10.3390/BRAINSCI14100982.

48. Signoret-Genest, J., Schukraft, N., L. Reis, S., Segebarth, D., Deisseroth, K., and Tovote, P. (2023). Integrated cardio-behavioral responses to threat define defensive states. Nat. Neurosci., 1–11. 10.1038/s41593-022-01252-w.

49. Koutsikou, S., Crook, J.J., Earl, E. V, Leith, J.L., Watson, T.C., Lumb, B.M., and Apps, R. (2014). Neural substrates underlying fear-evoked freezing: the periaqueductal grey–cerebellar link. J. Physiol. 592, 2197–2213. 10.1113/jphysiol.2013.268714.

50. Shanley, M.R., Miura, Y., Guevara, C.A., Onoichenco, A., Kore, R., Ustundag, E., Darwish, R., Renzoni, L., Urbaez, A., Blicker, E., et al. (2023). Estrous Cycle Mediates Midbrain Neuron Excitability Altering Social Behavior upon Stress. The Journal of Neuroscience 43, 736. 10.1523/JNEUROSCI.1504-22.2022.

51. Frontera, J.L., Aissa, H.B., Sala, R.W., Mailhes-Hamon, C., Georgescu, I.A., Léna, C., and Popa, D. (2020). Bidirectional control of fear memories by cerebellar neurons projecting to the ventrolateral periaqueductal grey. Nat. Commun. 11, 1–17. 10.1038/s41467-020-18953-0.

52. Popa, D., Spolidoro, M., Proville, R.D., Guyon, N., Belliveau, L., and Léna, C. (2013). Functional Role of the Cerebellum in Gamma-Band Synchronization of the Sensory and Motor Cortices. Journal of Neuroscience 33, 6552–6556. 10.1523/JNEUROSCI.5521-12.2013.

53. Sengupta, A., Bocchio, M., Bannerman, D.M., Sharp, T., and Capogna, M. (2017). Control of Amygdala Circuits by 5-HT Neurons via 5-HT and Glutamate Cotransmission. J. Neurosci. 37, 1785–1796. 10.1523/JNEUROSCI.2238-16.2016.

54. Herry, C., and Johansen, J.P. (2014). Encoding of fear learning and memory in distributed neuronal circuits. Nat. Neurosci. 17, 1644–1654. 10.1038/NN.3869.

55. Proville, R.D., Spolidoro, M., Guyon, N., Dugué, G.P., Selimi, F., Isope, P., Popa, D., and Léna, C. (2014). Cerebellum involvement in cortical sensorimotor circuits for the control of voluntary movements. Nat. Neurosci. 17, 1233–1239. 10.1038/nn.3773.

56. Kelly, K.J., Donner, N.C., Hale, M.W., and Lowry, C.A. (2011). Swim stress activates serotonergic and nonserotonergic neurons in specific subdivisions of the rat dorsal raphe nucleus in a temperature-dependent manner. Neuroscience 197, 251–268. 10.1016/J.NEUROSCIENCE.2011.09.011.

57. Locke, T.M., Soden, M.E., Miller, S.M., Hunker, A., Knakal, C., Licholai, J.A., Dhillon, K.S., Keene, C.D., Zweifel, L.S., and Carlson, E.S. (2018). Dopamine D1 Receptor–Positive Neurons in the Lateral Nucleus of the Cerebellum Contribute to Cognitive Behavior. Biol. Psychiatry 84, 401–412. 10.1016/J.BIOPSYCH.2018.01.019.

58. Chung, J.E., Magland, J.F., Barnett, A.H., Tolosa, V.M., Tooker, A.C., Lee, K.Y., Shah, K.G., Felix, S.H., Frank, L.M., and Greengard, L.F. (2017). A fully automated approach to spike sorting. Neuron 95, 1381–1394.e6. 10.1016/j.neuron.2017.08.030.

59. Carlson, E.S., Hunker, A.C., Sandberg, S.G., Locke, T.M., Geller, J.M., Schindler, A.G., Thomas, S.A., Darvas, M., Phillips, P.E.M., and Zweifel, L.S. (2021). Catecholaminergic Innervation of the Lateral Nucleus of the Cerebellum Modulates Cognitive Behaviors. Journal of Neuroscience 41, 3512–3530. 10.1523/JNEUROSCI.2406-20.2021.

60. Buccino, A.P., Hurwitz, C.L., Garcia, S., Magland, J., Siegle, J.H., Hurwitz, R., and Hennig, M.H. (2020). SpikeInterface, a unified framework for spike sorting. Preprint, 10.7554/eLife.61834 https://doi.org/10.7554/eLife.61834.

61. Ikai, Y., Takada, M., Shinonaga, Y., and Mizuno, N. (1992). Dopaminergic and non-dopaminergic neurons in the ventral tegmental area of the rat project, respectively, to the cerebellar cortex and deep cerebellar nuclei. Neuroscience 51, 719–728. 10.1016/0306-4522(92)90310-X.

62. Cutando, L., Puighermanal, E., Castell, L., Tarot, P., Belle, M., Bertaso, F., Arango-Lievano, M., Ango, F., Rubinstein, M., Quintana, A., et al. (2022). Cerebellar dopamine D2 receptors regulate social behaviors. Nature Neuroscience 2022 25:7 25, 900–911. 10.1038/s41593-022-01092-8.

63. Yu, S.K., Jung, H.S., Hall, F.S., and Linden, D.J. (2009). Dopamine Signaling Is Required for Depolarization-Induced Slow Current in Cerebellar Purkinje Cells. Journal of Neuroscience 29, 8530–8538. 10.1523/JNEUROSCI.0468-09.2009.

64. Saitow, F., Arakawa, R., and Suzuki, H. (2025). Cross-talk between dopamine and noradrenaline modulates glutamatergic transmission in the deep cerebellar nuclei. Sci. Rep. 15, 1–14. 10.1038/S41598-025-17213-9;SUBJMETA.

65. Novello, M., Bosman, L.W.J., and De Zeeuw, C.I. (2022). A Systematic Review of Direct Outputs from the Cerebellum to the Brainstem and Diencephalon in Mammals. The Cerebellum. 10.1007/s12311-022-01499-w.

66. Sacchetti, B., Baldi, E., Ambrogi Lorenzini, C., and Bucherelli, C. (2002). Cerebellar role in fear-conditioning consolidation. Proc. Natl. Acad. Sci. U. S. A. 99, 8406. 10.1073/PNAS.112660399.

